# The neural orchestra of aggression: neurogenetic network mapping of human aggressiveness

**DOI:** 10.1101/2025.10.31.685919

**Authors:** Jules R. Dugré, Macià Buades-Rotger, Stéphane De Brito

## Abstract

Over the last century, researchers have successfully mapped the core neural circuitry underlying aggression in non-human animals. In contrast, advances in human neuroimaging have been hindered by persistent challenges with reproducibility. Here, we adopted a recently developed network-based framework to test the hypothesis that seemingly heterogeneous findings in aggression research converge on a common brain network. We conducted network mapping to integrate functional and structural imaging findings of human aggression across 39 and 31 samples, respectively, revealing substantial overlap across both functional (up to 84%) and structural (up to 74%) imaging modalities. Strikingly, we found that these networks were largely explained by the expression of genes implicated in genome-wide association studies of aggression and related phenotypes. By integrating network-based approaches of neuroimaging data with gene expression, our work provides a reliable and comprehensive account of the neurogenetic architecture underlying aggressive behavior, resolving longstanding discrepancies in its neurobiology.

## Introduction

Aggression, often defined as a behavior intended to harm others ^1^, constitutes a major societal concern, affecting millions of lives and leading to significant social, economic and personal harm^2^. Decades of research spanning the past century have led in the identification of a brain circuit in non-human animals that constitutes the core aggression network ^3^. More precisely, the medial nucleus of the amygdala, ventromedial hypothalamus, and periaqueductal grey (PAG) are central to initiating aggressive responses to perceived threat across many species ^4, 5^. In human, both functional and structural neuroimaging studies in populations at risk for aggression ^6–10^ have documented the importance of additional subcortical (e.g., striatum, thalamus) and cortical regions (e.g., anterior insula, dorsomedial and lateral PFC) in populations at risk for aggression. More precisely, findings from a recent meta-analysis of functional magnetic resonance imaging (fMRI) studies on trait aggression supported involvement of the centromedial amygdala, but also highlighted many other brain regions known to be involved in executive function (i.e., intraparietal sulcus, angular gyrus) and social cognition (i.e., precuneus, middle temporal gyrus;^11^. In turn, the most recent meta-analysis of voxel-based morphometry (VBM) studies on trait aggression revealed a significant reduction of grey matter volume (GMV) in the medial prefrontal cortex ^6^. Taken together, these recent findings highlight that the neural circuitry of human aggression may involve a more widely distributed network than that of non-human animals.

### Shifting paradigm in Neuroimaging Research

Neuroimaging research is currently facing a replication crisis, which has been partly attributable to the poor test-retest reliability of commonly used fMRI tasks ^12^, scan length ^13^, sample size ^14,15^, and inconsistencies between studies ^16–18^. Indeed, neuroimaging meta-analyses of both fMRI ^7, 9, 11, 19–21^ and VBM ^6–10^ studies have reported notable spatial discrepancies, even when the meta-analyses include largely overlapping sets of studies. One possible explanation for such inconsistencies is that meta-analyses may be biased by the results of individual studies. Indeed, it has been shown that traditional coordinate-based meta-analyses are frequently driven by a few studies ^22–25^. Consequently, neuroimaging meta-analyses are highly sensitive to study selection criteria—a limitation exacerbated by substantial between-study variability (e.g., sample characteristics, methodology, etc.). A complementary explanation for the lack of robust meta-analytic findings is that traditional approaches rely on an erroneous assumption that brain regions operate as isolated units, yielding unstable estimates at a regional level. In contrast to this view, recent work indicates that complex behaviors can only arise from information propagation over distributed brain networks ^26–28^. As analogy, brain organization may be best conceptualized as a complex orchestra—organized into modules of highly interconnected units (e.g., strings, woodwinds) performing in harmony, rather than a group of soloists.

The localizationist view has historically dominated affective neuroscience, with early work seeking to identify focal brain regions underlying aggressive behavior. This line of work was started with early phrenological theories (e.g., mid-temporal regions, ^29^), followed by surgical brain resections (e.g., temporal lobe, Burghardt cited in ^30^), and lesion studies (e.g., ventromedial PFC, ^31^), through to modern brain stimulation studies in rodents (e.g., hypothalamic attack area, ^32^) and humans (e.g., posterior hypothalamus, ^33^). These distinct yet interrelated approaches have produced inconsistent results, calling into question whether human aggression can truly be localized to a few specific brain regions. In contrast, contemporary models in human neuroscience emphasize the dynamic harmonization of signal fluctuations across a distributed set of interconnected regions to produce (mal)adaptive behavior ^27^. This allows researchers to test alternative hypotheses, such as *neurobiological equifinality* ^22, 28^, in which different neurobiological pathways can lead to the same outcome (e.g., aggressive behavior). Supporting this, recent meta-analyses have shown that functional connectivity between heterogeneous brain regions can capture neural commonalities between studies, beyond clinical and methodological differences ^24, 34–37^. Indeed, recent meta-analytic advances have demonstrated that shifting from a localizationist approach to a network-based model can indeed substantially increase replicability among functional ^22^ and structural ^23^ imaging studies in adult with psychopathy. Despite recent evidence suggesting that studies on aggression may share a common network ^38^, the biological mechanisms explaining this association remain poorly understood.

### Bridging levels of understanding in aggression research

Other research fields, like genomics, can also critically contribute to our understanding of the neurobiological basis of aggression. Genome-wide association studies (GWAS) have identified hundreds of potential genetic variants associated with aggression and antisocial behavior ^39–43^. Yet, the specific contributions of these genes to brain structure and function (e.g., synaptic signaling, neuronal projection, neurogenesis) remain poorly understood. Recent methodological advancements now enable the co-localization of neuroimaging findings with gene expression patterns in the brain ^44, 45^. For example, Rokicki and colleagues ^46^ recently revealed that GWAS-derived single nucleotide polymorphisms related to antisocial behaviors ^47^ were highly expressed in cerebellar, subcortical, and prefrontal regions, particularly in the anterior cingulate cortex and orbitofrontal cortex. Yet, we do not know whether gene expression in the brain aligns with differences in neural activity and morphology in individuals with varying levels of aggressive behavior. Uncovering the links between genetic, neurostructural, and neurofunctional data is thus crucial for establishing a comprehensive and multi-level neurobiological framework that can facilitate the prognosis and treatment of aggression.

Here, we re-analyzed data from our recently published functional and structural neuroimaging meta-analyses on aggression ^11, 48^. First, we aimed to investigate the spatial convergence among peak coordinates between studies as a measure of robustness of prior findings. We hypothesized weak spatial overlap between studies, in line with previous work indicating that prior meta-analytic findings may be driven by few studies ^23, 37, 49, 50^. Second, we investigated whether these spatially heterogeneous brain regions across aggression literature may map onto a common brain network. Consistent with the hypotheses proposed in previous studies ^22, 23^, we hypothesized that neuroimaging studies on aggression would show greater replicability when using a network-based approach achieving a substantial overlap across studies (>60%). Finally, for the first time, we aimed to identify whether genes identified in genome-wide association studies (GWAS) on aggression and related phenotypes were expressed in similar regions to those of aggression-related brain networks. Across all analyses, we tested for differences between reactive (impulsive, “hot-blooded”) and proactive (instrumental, “cold-blooded”) aggression, which show partial yet distinct, neurobiological mechanisms ^48, 51, 52^. By doing so, we integrate data across multiple levels of analysis to provide a comprehensive account of the neurogenetic architecture underlying aggressive behavior.

## Methods

### Literature search

A systematic search using three search engines (Google Scholar, PubMed and Web of Science) was conducted up to May 1st, 2023 as described in ^6, 11^. The following search terms were used: (aggress* OR violen*) AND (neuroimaging OR fMRI OR VBM OR functional neuroimaging OR structural neuroimaging OR task-based OR voxel-based). Irrelevant records and duplicates were first excluded. Full texts of the resulting studies were subsequently screened. An additional search was conducted by cross-referencing the reference lists of the included articles.

### Study selection

Articles were included if they met the following criteria: (1) original study published in a peer-reviewed journal in English; (2) inclusion of a validated measure of aggressive behavior (i.e., questionnaires, interviews); (3) inclusion of grey matter volume and/or task-based fMRI modality; (4) conducted a group-comparison or whole-brain regression assessing the dimensional relationship between voxels and the severity of trait aggression (5); reported peak coordinates and significant effects across the whole-brain (i.e., voxel-wise) [null or peak coordinates]. We excluded studies assessing aggressive behaviors within the scanner without a validated measure of aggression (e.g., severity of a noise blast). These were excluded because of the poor correlation between laboratory tasks with trait aggression questionnaires (see recently ^53, 54^) and because the neural correlates of laboratory-based aggression have been extensively analyzed ^20, 21, 55^. In addition, we excluded studies reporting only one-sample t-tests. This exclusion criterion was chosen given that we aimed to examine the inter-individual differences in trait aggression and brain functions and structures. Preferred Reporting Items for Systematic Reviews and Meta-Analyses (PRISMA) was followed across the meta-analysis steps (see Supplementary Tables 1-2)

As described in recent neuroimaging meta-analyses of functional and structural studies ^6,11^, a total of 67 fMRI and 36 VBM studies were included (Please refer to Supplementary Tables 3-4). More precisely, general aggression was measured across 35 fMRI (42 samples) and 26 VBM (32 samples) studies. From these, 39 fMRI and 31 VBM samples were included in the current meta-analysis after removing samples with null findings. As we aimed to test generalizability across subtypes of aggression, studies measuring reactive and proactive aggression were also included. After removing samples with null findings, the current meta-analysis included 37 fMRI and 20 VBM samples on reactive aggression, and 19 fMRI and 11 VBM samples on proactive aggression.

### Normative network mapping

We applied a normative network mapping approach to identify whether heterogeneous peak coordinates reported across studies on aggression may be linked to a common network ^24, 34, 35, 37, 56, 57^. Briefly, a 4-mm sphere was created around each coordinate from each sample to create a sample-level binary mask. Then, we generated the network profile of each sample-level mask using both functional ^22^ and structural ^23^ neuroimaging data from 1,000 healthy subjects (ages 18 to 35 years old, 50% females) of the Brain Genomics Superstruct Project ^58, 59^. Details about the preprocessing can be found elsewhere (https://doi.org/10.7910/DVN/ILXIKS). For task-based samples, we first extracted the time-course of voxels within each sample-level mask (4mm spheres) and correlated them to the time course of every other voxel in the brain for each of the 1,000 healthy participants, yielding a sample-level map per subject (*n* sample x 1,000 participants). For VBM sample, we extracted the mean volume of each sample-level mask (4mm spheres) in each of the 1,000 healthy individuals and correlated them to the mean volume of each voxel in the brain, yielding a similar sample-level map per subject (*n* studies x 1,000 participants). This was done using partial correlation with Total Intracranial Volume (TIV) as covariate to control for inter-individual differences in brain size. Then, these 1,000 sample-level maps were combined in a voxel-wise one-sample *t*-test, resulting in a single sample-level map representing the group-level connectivity profile of peaks of that study.

To ascertain if heterogeneous peaks do indeed map onto a common network, two analyses were conducted by quantifying: 1) the average network strength across samples; and 2) the degree of overlap across samples. This approach allowed us to identify a group-level network, but also to determine how many studies converge on that network. For the first analysis, sample-level maps were converted to Fisher’s z scores and combined in a voxel-wise one-sample *t*-test to identify a network that is more consistent than expected by chance ^60^. Statistical significance was set using cluster-based Threshold-Free Cluster Enhancement (TFCE) and Family-Wise Error corrections (FWE TFCE<0.05, 5,000 permutations). For the second analysis, aiming to quantify the degree of overlap across samples, T map of each sample was thresholded (T>5) and binarized. We then summed these binarized t-maps to quantify, for each voxel, the percentage of studies sharing the same overarching network, despite reporting heterogeneous peak locations. This threshold has been chosen according to previous studies using a similar approach ^22, 23, 34^. Note also that the use of a fixed and stringent threshold circumvents the inherent variability of other methods such as False Discovery Rate correction ^61^ between samples. These analyses were carried out for task-based samples (functional connectivity) and VBM studies (structural covariance), separately. Moreover, although the main analyses report findings across measures assessing aggression in general, analyses were re-analyzed across independent samples on subtypes of aggression, namely reactive and proactive aggression. As previously described, aggression subtypes were classified as either reactive or proactive based on the questionnaire and specific subscale(s) used ^6, 11^.

#### Contribution of intrinsic connectivity networks

Intrinsic connectivity is fundamental to our understanding of signal propagation across the brain. Consequently, we examined whether our findings may have been driven by connectivity within versus between particular networks. To do so, we used 8 intrinsic connectivity networks, namely the Schaefer-400 parcels 7 Networks ^62^ and a subcortical network ^63^. Effect sizes (Cohen’s *d*) were computed by the difference between WITHIN_tval_ (i.e., mean t-value within a network) and BETWEEN_tval_ (i.e., mean t-value outside the given network) divided by the pooled standard deviation. This measure indexes the degree to which the observed results reflect intra- vs inter-network coupling.

### Genetic contribution to aggression-related brain networks

#### Neural mapping of gene expression

We also investigated which genes were predominantly expressed in aggression-derived functional and structural brain networks. To do so, we first extracted single nucleotide polymorphisms (SNPs) identifiers across 21 GWAS (26 samples, see Table 1) on aggression and related phenotypes (e.g., Anger/Irritability, Conduct Disorder/Problems, Antisocial Personality) derived from recent systematic reviews and meta-analyses ^39–43^. Only SNPs that met the conventional threshold of p<1×10^−5^ were included in subsequent steps. Second, following recent approach ^46^, the Variant-to-Gene (v2g) tool from the Open Targets Genetics was used to map SNPs to genes, using multiple types of evidence (i.e., including positional information, chromatin interaction data, expression/protein/splicing quantitative trait loci [QTL], and in silico functional predictions ^64^). This resulted in a total of 481 mapped genes across 26 samples, from which 458 were non-overlapping.

**Table 1.** Genome-Wide Association Studies and Meta-analyses on Aggression and Related Phenotypes.

| First Author, Year | Sample Characteristics |  |  | Genetic Markers |  |
| --- | --- | --- | --- | --- | --- |
|  | N<sup>A</sup> | Age Range<sup>B</sup> | Phenotype | N of Mapped Genes | N of Stable Genes (DS > 0.1) |
| Aebi, 2016<sup>75</sup> | 750 | 5–18 | Oppositionality | 11 | 5 |
| Anney, 2008<sup>76</sup> | 938 | 5–17 | Conduct Problems (ADHD) | 25 | 14 |
| Bani-Fatemi, 2020<sup>77</sup> | 205 | 18–75 | Aggression (Schizophrenia) | 4 | 2 |
| Brevik, 2016*<sup>78</sup> | 1,060 | 17–75 | ODD | 19 | 8 |
| Demontis, 2021*<sup>79</sup> | 3802 | NA | ADHD+DBD | 39 | 25 |
| Dick, 2011<sup>80</sup> | 3963 | 18–74 | Conduct Problems/Disorder | 20 | 10 |
| Ip, 2021*<sup>81</sup> | 87 485 | 1.5–18 | Aggression | 43 | 23 |
| Li et al., 2023*<sup>82</sup> | 644, 1081, 1029, 463 | 17-75 | ASPD Count | 61 | 37 |
| McGue, 2013<sup>83</sup> | 7,188 | 16.5-21 | Conduct Problems | 7 | 4 |
| Merjonen, 2011<sup>84</sup> | 987 to 1776 | 15-45 | Irritability | 7 | 1 |
| Mick, 2011<sup>85</sup> | 341 | M=10.8 | Dysregulation | 5 | 3 |
| Mick, 2014<sup>86</sup> | 845 & 515 | 45–64 | Anger (Reaction+Temper) | 13 | 4 |
| Montalvo-Ortiz, 2018<sup>87</sup> | 2185 & 1,362 | M=40.67 & 36.47 | Aggression (Cannabis Use) | 89 | 42 |
| Pappa, 2016*<sup>88</sup> | 18,988 | 3-15 | Aggressive behaviors | 5 | 4 |
| Rautiainen, 2016<sup>89</sup> | 370 | M=34.5 | ASPD | 11 | 3 |
| Salvatore, 2015<sup>90</sup> | 1379 | 18–79 | ASPD Criteria | 8 | 1 |
| Sonuga-Barke, 2008<sup>91</sup> | 909 | 5–17 | Conduct Disorder (ADHD) | 10 | 5 |
| Tielbeek, 2012<sup>92</sup> | 4816 | 24-41 | ASPD | 8 | 4 |
| Tielbeek, 2017*<sup>93</sup> | 16,400 | 4-41 | Antisocial Behaviors | 49 | 26 |
| Tielbeek, 2022*<sup>47</sup> | 85,359 | 11-85 | Antisocial Behaviors | 38 | 17 |
| Tiihonen, 2015<sup>94</sup> | 360 & 84 | M=37.4 & 36.7 | Violent Offending | 9 | 8 |
| **Non-Overlapping** | - | - | - | 502 | 246 |
| **Total** | - | - | - | 481 | 230 |

Note. A = Multiple sample sizes correspond to different samples within the study, B = Age range is displayed, and mean age of the sample when the age range was not available. \* Meta-analysis of multiple samples. ADHD = Attention-Deficit/Hyperactivity Disorder; ODD = Oppositional Defiant Disorder; DBD = Disruptive Behavior Disorder; ASPD = Antisocial Personality Disorder.

Regional microarray gene expression data were obtained from six post-mortem brain provided by the Allen Human Brain Atlas ^45^. Each donor brain was sampled in 363–946 locations, either in the left hemisphere only (*n* = 6), or over both hemispheres (*n* = 2). Data processing was conducted following Rokicki and colleagues ^46^ and involved computing one high-density (interpolated using 10-nearest neighboring tissue samples at each voxel and bidirectional mirroring) and one atlas-based map per gene (Schaefer-7 400 parcels, ^62^ with the python toolbox abagen ^65^. Data was normalized using the default scaled robust sigmoid method ^44^. Following a recent study, we only selected genes with a differential stability above 0.1 ^66^. This ensures a minimal consistency in cortical expression across the six donors and reduces the likelihood of chance findings ^67^. From the 458 genes, 148 had a differential stability of less than 0.1 and were further removed. From the remaining 83 genes, 3 had different aliases from the Open Targets Genetics, while 80 were not found in the Allen Human Brain Atlas, leaving a final dataset of 230 gene expression maps.

#### Joint variation between functional, structural, and gene expression maps

To clarify the genetic contribution to brain networks underpinning aggression, a Partial Least Squares (PLS) analysis was conducted to link GWAS-derived genes and brain networks (i.e., function and structure). This approach extracts latent variables that maximize the covariance between modalities which, in this case, are gene expression maps and aggression-related structural and functional networks. Importantly, PLS allows to identify which components within each latent variable contribute most to the observed association ^68^. Here, we only focused on the first latent variable accounting for most variance between gene expression and brain networks. Because spatial auto-correlation might inflate the estimated association between brain maps ^69, 70^, we investigated the correlation between gene expression and brain networks using distance-dependent cross-validation ^66, 71^. Briefly, for each region of the Schaefer-400 atlas (source node), we selected the 75% closest regions in surface space as the training set, leaving the remaining 25% of regions for test set. The model was first fitted on the training set, and weights were projected onto the test set. The correlation between gene expression and brain networks in the test set (mean out-of-sample correlation) was compared against a permuted null model (1,000 repetitions) constructed by repeating the cross-validation on permutations of the brain networks matrix, after spherical projection, random rotation, and reassignment ^71^. To better interpret these findings, weights attributed to each of the 230 genes were extracted, and studies were ranked based on their contribution to this latent variable (i.e., sum of weights divided by the number of genes). Genes whose brain expression contributed positively to the aggression brain network were analyzed using gene-set enrichment to identify enriched biological processes (GO Biological Process 2025, ^72^) via Enrichr ^73^.

We subsequently conducted supplemental analyses to test the robustness of our findings. First, we tested whether the PLS findings might be impacted by inter-subject variance of gene expression across donors, namely, differential stability. To do so, we computed the Spearman correlation coefficient between absolute values of attributed weights for each gene and the differential stability score to ensure that the most important genes in the explanation of aggression brain networks are robust across donors. Second, given that the PLS was conducted only on cortical expression, we also examined whether the gene-imaging modality latent variable was generalizable across subcortical regions. This was done by applying the same weights identified in the PLS analysis to gene expression and imaging modalities across 14 subcortical regions ^74^. Statistical significance was assessed via permutation testing by shuffling the subcortical regions 5,000 times. We finally explored whether the PLS model generalized across independent samples which assessed reactive and proactive subtypes of aggression.

## Results

### Normative network mapping

To assess spatial heterogeneity in the aggression literature, peak coordinates from functional and structural imaging studies were overlapped, with each peak modeled as a 4 mm sphere. Findings revealed low convergence across the 39 independent samples derived from 35 functional neuroimaging studies, as evidenced by peak overlap reaching a maximum of 10.25% (4 out of 39 samples)(Fig. 2). Across the 31 independent samples derived from 26 VBM studies, the maximal spatial overlap between peak locations was 9.67% (3 out of 31 samples), similar to what was observed across fMRI studies (Fig. 2).

**Fig 1.**
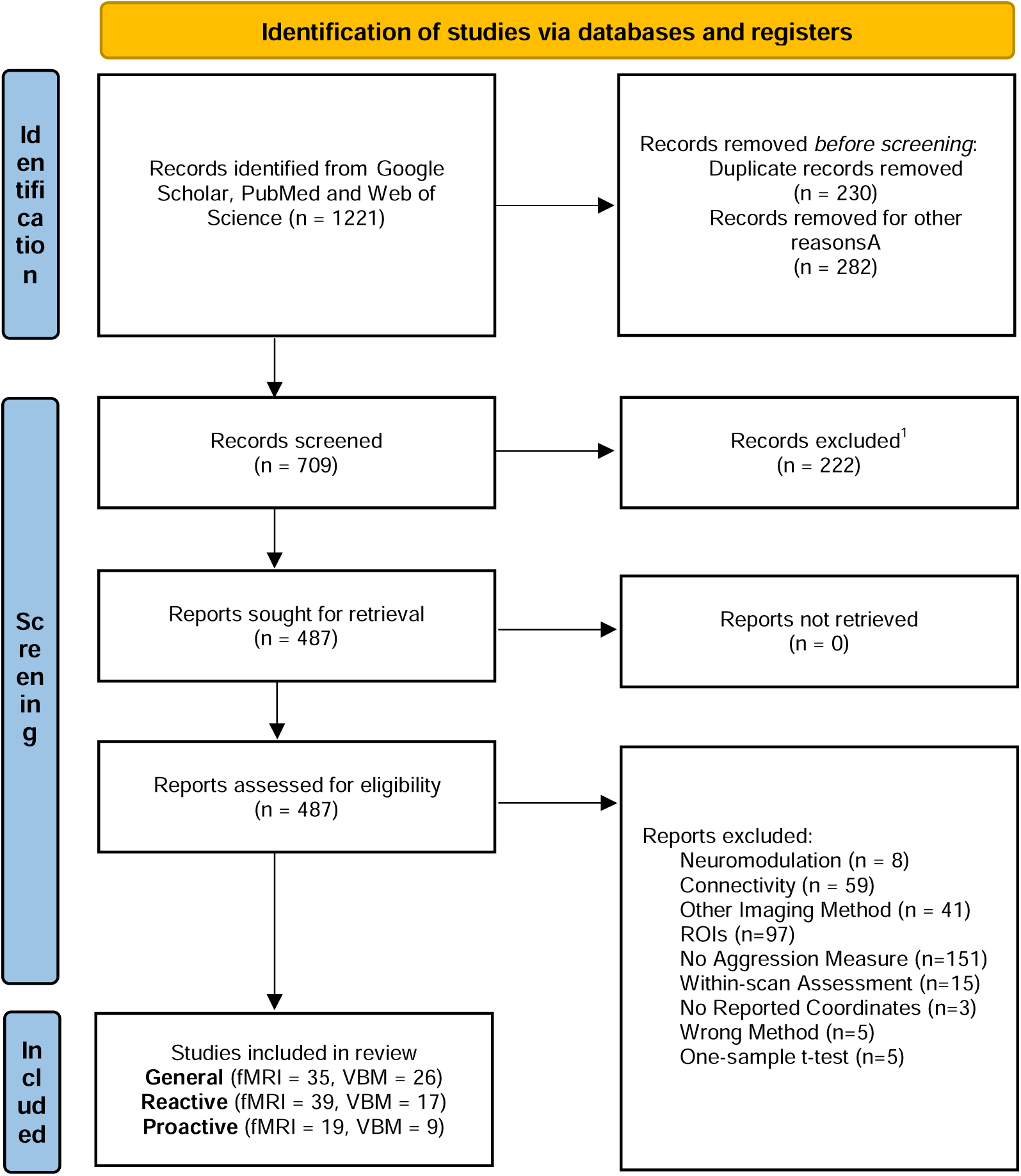
PRISMA (Preferred Reporting Items for Systematic reviews and Meta-Analyses) Flowchart. ^1^ Reasons for exclusion were Literature Reviews (n=138), Animal Studies (n=19), Absence of fMRI or sMRI data (n=26), Case Studies (n=6), Abstract, Book Chapter, and Thesis (n=33). ROIs = Region-of-Interest; fMRI = Functional Magnetic Resonance Imaging; VBM = Voxel-Based Morphometry; General: general aggression; Reactive: reactive (also termed impulsive or “hot-blooded”) aggression; Proactive: proactive (also termed instrumental or “cold-blooded” aggression).

**Fig. 2.**
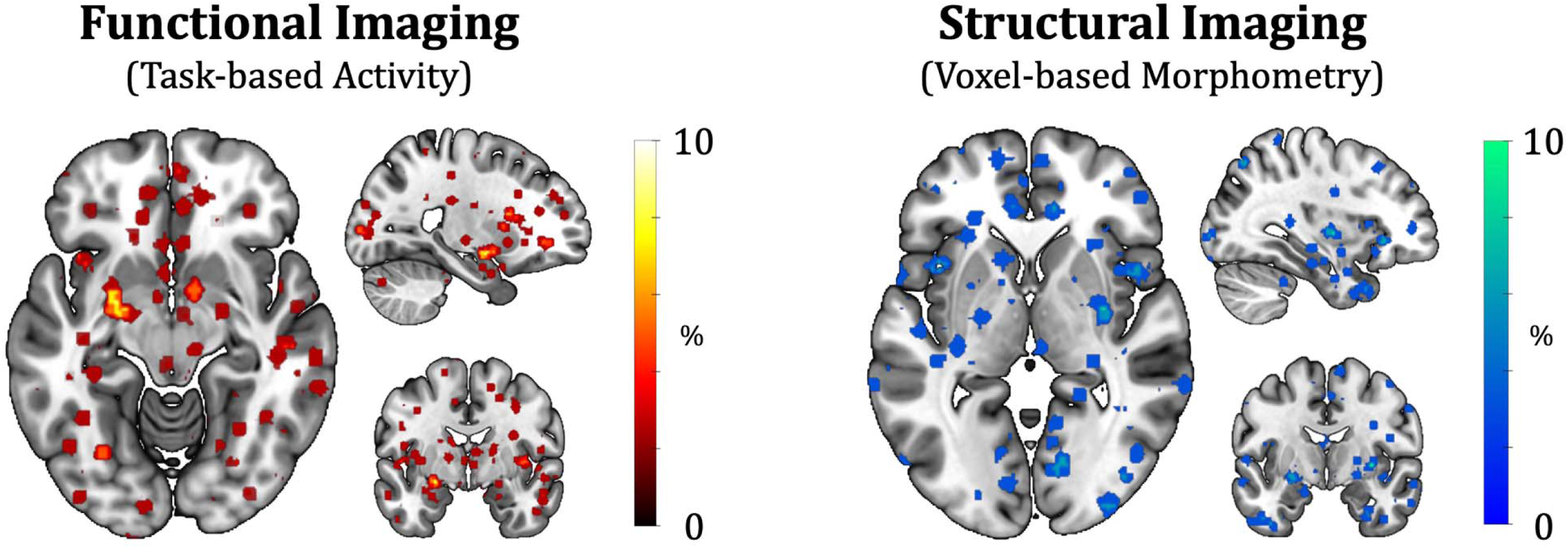
Low reproducibility among neuroimaging studies on trait aggression. Spatial overlap in peak coordinates across neuroimaging samples revealed that, at most, 4 out of 39 functional imaging samples converged on the same peak locations (10.25%; left panel), and only 3 out of 31 structural imaging samples showed overlapping peaks (9.67%; right panel). Peak coordinates were modeled using a 4 mm³ sphere.

Conducting network mapping across functional neuroimaging samples unveiled a statistical significant brain network underpinning task-based activation samples on aggression (pFWE-tfce<0.05, Fig. 3A), which was mainly characterized by stronger connectivity from nodes of the subcortex (*d* = 1.43) and the default-mode (*d* = 1.10), and frontoparietal networks (*d* = 0.57) to a lesser extent (Fig. 3A). More precisely, this network reached up to 84.62% overlap across samples, particularly in the caudate nucleus extending into the bed nucleus of the stria terminalis (Fig. 3A, Supplementary Table 5). Other replicable brain regions included the perigenual anterior cingulate cortex (74.36%), left and right ventral anterior insula (both 74.36%), thalamus (74.36%), left lateral amygdala extending to the anterior hippocampus (66.67%), and left and right mammillary tracts extending to the posterior hypothalamus (71.80%, and 74.36% respectively). The replicability and relative importance of the subcortex and default mode network were comparable across independent samples for both reactive (37 samples) and proactive (19 samples) aggression (Supplementary Fig. 1B).

**Fig 3.**
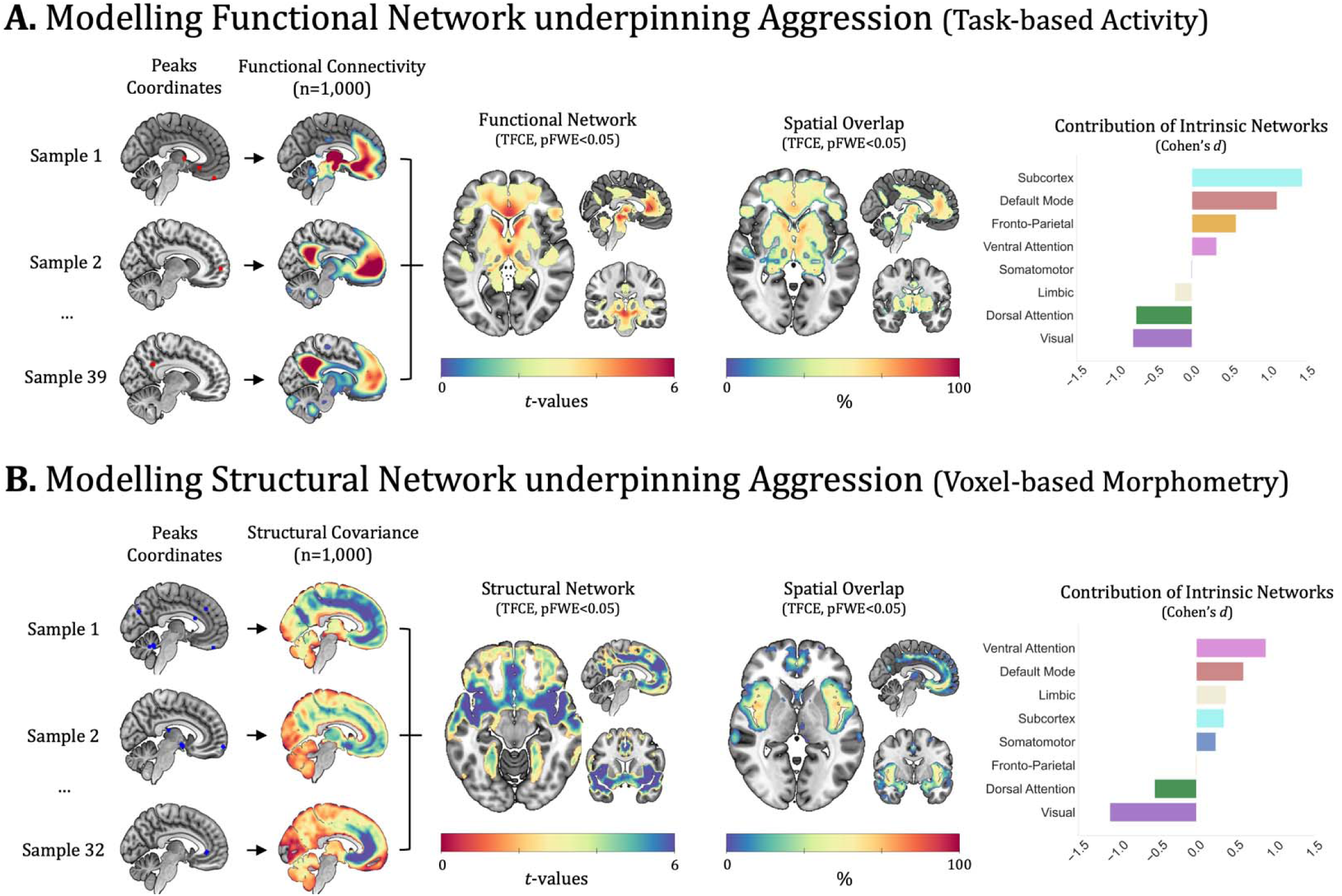
Modelling the Functional and Structural Brain Networks of Trait Aggression. Panel. **A.** Peak coordinates reported across neuroimaging studies on trait aggression were modeled using a 4 mm spheres. Within each functional imaging study (upper row), a network profile of each coordinate was generated and aggregated using resting-state functional connectivity, while structural covariance of grey matter volume was used for generating a network profile for each peak coordinate within each structural neuroimaging study (**Panel B**). These study-level network profiles were then fed into a one-sample permutation tests to generate a group-level network that is more consistent than expected by chance (Family-Wise Error and Threshold-Free Cluster Enhancement corrected at p<0.05 with 5,000 permutations). Finally, we examined the contribution of 8 intrinsic networks (Schaefer-7 and subcortex) to the modular organization of the functional **(A)** and structural **(B)** brain networks associated with aggression. Values of the bar chart represent the importance of each intrinsic network, as calculated by Cohen’s *d*.

Structural network mapping revealed a statistically significant brain network underpinning VBM samples on aggression (pFWE-tfce<0.05, Fig. 3B). This network was mainly characterized by stronger structural covariance of nodes from the ventral attention/salience (*d* = 0.88) and default-mode networks (*d* = 0.60), followed by limbic (*d* = 0.38) and subcortical networks (*d* = 0.35; Fig. 3B). More precisely, revealed a maximum overlap of 74.19% located in the dorsal aspect of the right middle insula (Fig. 3B., Supplementary Table 6). Other replicable brain regions included the left and right superior temporal gyri extending to the posterior insula (70.97%, and 67.74%, respectively), the dorsal part of the left middle insula (67.74%), the orbitofrontal cortex (67.74%), the right posterior insula (67.74%), the left and right claustrum extending to anterior insula (67.74%, and 64.5%, respectively), and the bilateral temporal pole extending to superficial amygdala (both 61.29%). This replicability as well as the importance of the replicability and relative importance of the ventral attention and default-mode network were also comparable across independent samples for both reactive (20 samples) and proactive (11 samples) aggression (see Supplementary Fig. 1B).

Notably, functional and structural network maps were moderately-to-highly correlated across cortical regions (*r* = 0.67, *P*_spin_ < 0.001) but not across subcortical regions (*r* = 0.03, *P*_shuffle_ > 0.05) (Supplementary Fig 2). A similar pattern was found for reactive aggression (cortical *r* = 0.63, *P*_spin_ < 0.001; subcortical *r* = 0.01, *P*_shuffle_ > 0.05), but not proactive aggression (cortical *r* = 0.17, *P*_spin_ > 0.05; subcortical *r* = 0.05, *P*_shuffle_ > 0.05) (Supplementary Fig 3), suggesting that the functional-structural overlap was more prominent for reactive than for proactive aggression.

### Genetic contribution to aggression brain networks

The 230 genes derived from the 21 GWAS on aggression and related phenotypes, identified a latent variable that linked the spatial organization of gene expression and aggression brain networks (*r* = 0.68, 46% of the covariance) (Fig 4A). The robustness of this model was measured using a distance-dependent cross-validation to assess the effect of spatial autocorrelation. The mean out-of-sample (top 25% farthest region to the source node) correlation between gene expression and brain network scores was *r* =.59 (*P*_spin_ = 0.012) (Fig 4B.). Precisely, the PLS assigned similar weights to both functional (w = 0.60) and structural (w = 0.79) networks, suggesting that the identified latent variable mostly reflected the shared neural patterns across imaging modalities. The set of 126 genes contributing positively to the explanation of aggression brain networks, were significantly associated with biological processes including cell-cell adhesion, synaptic membrane adhesion and regulation of neuron projection development (Supplementary Table 7). Exploring the weights assigned to each gene revealed that *SYT10* (synaptic vesicle release), *TENM3* (axon development and synaptic organization), *NAV3* (axonal growth), *SCN9A* (pain signaling and sensory neurons), and *ANKRD50* (endosome sorting), had greater contribution. Of the genes whose brain expression negatively correlated with aggression brain networks, *ZDHHC2*, *SLC17A6*, *SRXN1*, *KLF12*, and *SETD7* contributed most negatively (Fig. 4C).

**Fig 4.**
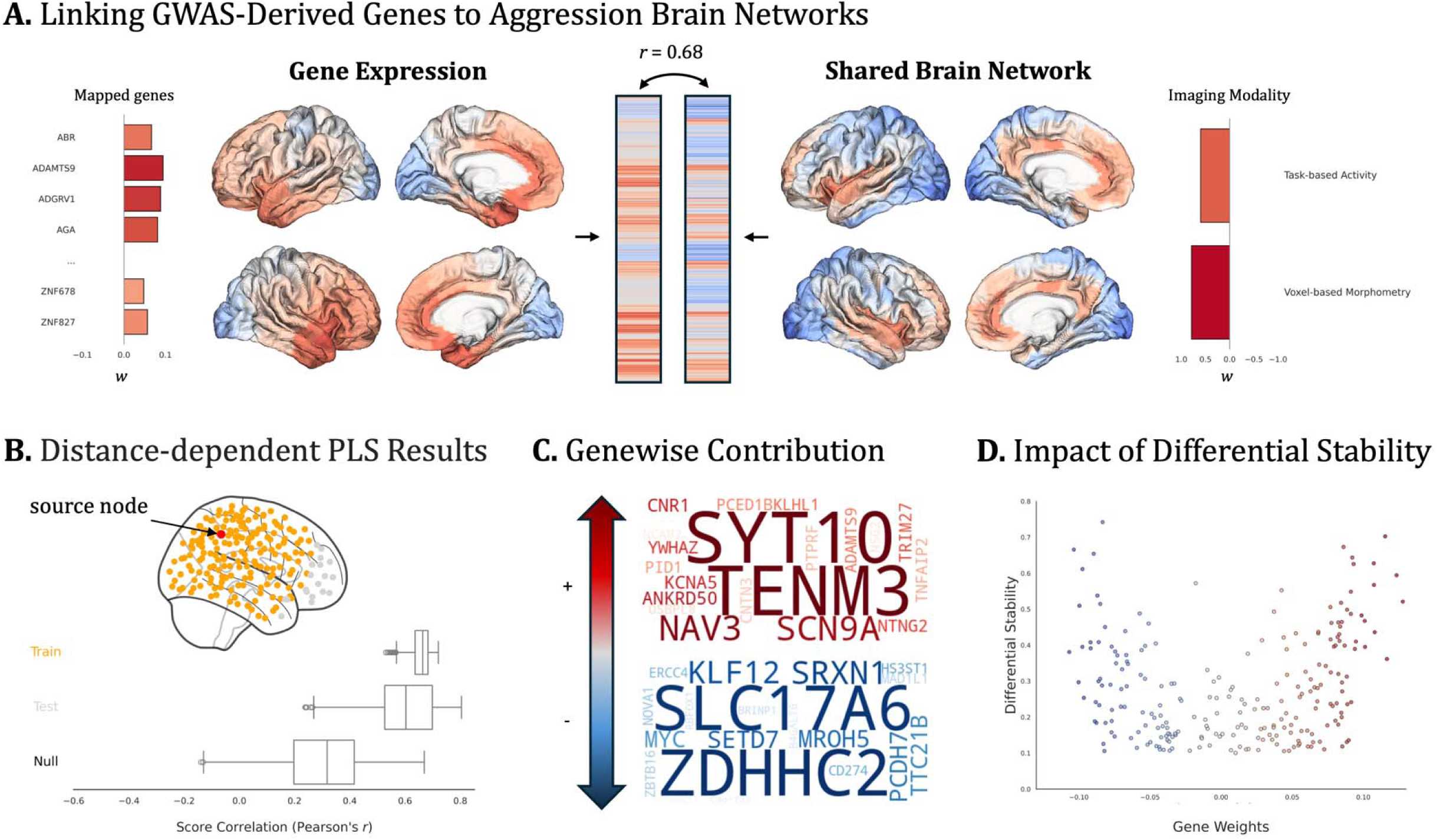
Understanding the genetic contributions to aggression brain networks. Panel. **A.** Depiction of the Partial Least Squares (PLS) analysis used to map the gene-brain connectivity interplay. **Panel B.** Results of the distance-dependent cross-validation conducted to minimize spatial autocorrelation.

Quality control subanalyses showed a strong correlation (ρ = 0.52, p < 1.00E-16) between PLS-weights assignment and differential stability (ρ = 0.55 p < 1.00E-10 for positive weights only & ρ = -0.47 p < 1.00E-06 for negative weights only), meaning that the most important genes were those with the lowest inter-subject variance in cortical gene expression across the six donors (Fig. 4D). Moreover, the latent variable identified from cortical brain regions also generalized across subcortical structures (*r* = 0.51, *P*_shuffle_ = 0.031, see Supplementary Fig 4B), but also across subtypes of aggression, in particular reactive aggression (cortical *r* = 0.64, *P*_spin_ = 0.0025; subcortical *r* = .62, *P*_shuffle_ = 0.0098) and only across cortical regions for proactive aggression (cortical *r* = .57, *P*_spin_ = 0.001; subcortex *r* = 0.43, *P*_shuffle_ = 0.064) (Supplementary Fig 5).

Briefly, each of the 400 Schaefer-7 cortical atlases were used as a source node (red point) to stratify data into training (yellow points, 75% closest regions) and test (grey points, 25% farthest) sets. To assess the statistical significance of the mean out-of-sample correlation (test set), a null distribution of the average correlation was constructed by randomly rotating the neuroimaging matrix before the PLS analysis. **Panel C.** shows the weights of the top 20 genes in each direction (positive, red; negative, blue) that explain the spatial distribution of brain networks underpinning aggression. **Panel D.** illustrates the strong correlation (ρ = 0.52, p < 1.00E-16 on absolute values) between differential stability of genes among the 6 donors and weights attributed to genes in the PLS (positive [red] & negative [blue] weights). This highlights that the most important genes in the explanation of aggression brain network are the most stable across donors, that is less inter-subject variance in cortical gene expression.

## Discussion

Despite substantial progress over the last decades, neuroimaging research on aggression has not converged in delineating the network of brain regions involved in human aggression. Here, we show that the seeming divergence across functional and structural neuroimaging studies share common neural architectures. Crucially, we show for the first time that these brain networks strongly overlap with the expression of genes previously associated with aggression and related clinical phenotypes (e.g., irritability, conduct disorder, antisocial personality disorder). Taken together, our findings demonstrate the usefulness of integrating multiple levels of analysis to understand the neurobiological architecture in human aggression.

### Brain networks associated with Aggression

Our findings align with the growing body of literature emphasizing the brain as a complex system of densely interconnected regions ^26–28^. Notably, peak locations of functional and structural imaging studies were highly heterogeneous, as demonstrated by their low spatial convergence. Nevertheless, by adopting a network-based approach that embraces inter-individual variability ^27, 28, 95^, we demonstrate how this apparent discrepancy across studies can be reconciled and potentially address the lack of reproducibility in neuroimaging research.

Normative network mapping of functional neuroimaging studies demonstrated the highest overlap in multiple subcortical structures previously linked with aggression such as the caudate nucleus, thalamus, hippocampus, and posterior hypothalamus ^4, 5^. Moreover, we observed that regions of the default mode network (mPFC/pgACC), and less prominently regions of the frontoparietal network, contributed the most to the characterization of the aggression-derived functional network. Similarly, previous meta-analyses demonstrated a conjunction between aggression-related brain activity and both mentalizing system and cognitive control areas ^9, 11^. Across species, the subcortex and midbrain play critical roles in detecting threats and orchestrating defensive responses ^4^. Cortically, the default mode network is involved in self-referential processes such as the understanding of others’ motives and intentions ^96^. Impairments in these processes are known risk factors to aggression ^97^, and may either contribute to the suppression of aggressive impulses or facilitate their strategic deployment ^98^. Concurring with recent lesion work ^99^, we highlighted the role lateral PFC in aggression, alongside other frontoparietal regions (e.g., inferior frontal gyrus, inferior parietal lobule), crucial for cognitive flexibility and successful inhibitory control ^100^. Intriguingly, this functional network shared several neurobiological similarities with the structural network, especially across the cortex (see Supplementary Material). These findings offer insight into how anatomical alterations may contribute to variations in brain function underlying aggression. Indeed, structural neuroimaging studies converged onto a network mainly formed by covariance of regions assigned to ventral attention (e.g., insula, claustrum, midcingulate cortex) and default mode networks (e.g., ventromedial prefrontal cortex/orbitofrontal cortex, superior temporal gyrus). Dovetailing with our findings, white matter integrity of a frontotemporal-subcortical network ^101^, and structural covariance between the superficial amygdala and anterior insula have previously been linked to aggression ^102^. Compromised integrity of the uncinate fasciculus has recently been identified as a core white matter tract in patients with brain lesions who subsequently committed violent crimes^103^. These findings suggest that anatomical pathways linking fronto-temporal and subcortical structures may underlie the shared deficits across functional and structural networks identified in this study, further reinforcing the importance of our network-based approach over traditional localizationist frameworks.

Overall, our findings are highly relevant for brain stimulation research aimed at improving clinical outcomes ^104^. For instance, the posterior hypothalamus, which demonstrated up to 74.36% replicability in the aggression-dependent functional brain network, has been recently identified as a promising deep-brain stimulation target ^33^. Intriguingly, the literature on transcranial direct current stimulation, which has so far focused on the mPFC, dlPFC, and vlPFC, has failed to show significant effects ^105^. In this context, our findings may help researchers optimize electrode placement (both anodal and cathodal) by avoiding configurations in which electrodes of opposing polarity are positioned within the same network, a factor thought to reduce stimulation efficacy. Furthermore, our findings may also provide further biological explanations about why cognitive-behavioral therapy ^106^, which typically involve a social perception module to identify and manage hostile attribution and attentional biases ^107^, as well as mentalizing-based treatment ^108^, produces efficacious results in reducing aggressive behaviors.

### Gene expression in aggression-related brain networks

One key challenge that remains is explaining how structural and functional brain networks may be linked, despite being driven by distinct regions/intrinsic networks. Here, we showed that most aggression-related genes derived from GWAS are integral to neurobiological processes that produce neurostructural alterations, with potential downstream effects on brain function. Indeed, we demonstrated that GWAS-related genes map onto a common structural network that significantly correlates with findings from VBM studies on trait aggression. To gain a more detailed understanding of which genes may drive this effect, partial least squares regression analysis identified a latent variable that accounted for 46% of the covariance between gene expression and imaging-derived brain networks. Importantly, we demonstrated that this effect generalized to subcortical structures as well as across reactive and proactive aggression—two subtypes that differ in their motives but predominantly align in terms of neurobiological correlates ^109^.

Many genes identified in GWAS on aggression and related phenotypes showed cortical expression that spatially resembled aggression brain networks. Interestingly, biological processes characterizing these genes reflected cell-cell adhesion, synaptic membrane adhesion, and regulation of neuron projection development. These processes highlight the importance of neural circuit formation and further argues in favor of adopting network-based approaches to understand the neurobiological architecture of aggression. Genes whose expression contributed most positively to the gene-brain covariance structure were more specifically implicated in synaptic vesicle release (*SYT10*), axon guidance (*TENM3*), and pain and sensory processing (*SCN9A*)—all processes relevant for emotional reactivity and behavioral regulation. Importantly, our results are solely based on genetic association studies, and possibly miss out on potential gene-environment interactions and other genomic processes that have been shown to be important for the development of aggression ^110, 111^. Nevertheless, our findings provide a foundation for targeted longitudinal studies designed to elucidate the genomic mechanisms underlying aggressive behavior.

## Limitations

There are several limitations that need to be acknowledged. First, neuroimaging studies on aggression vary significantly clinically (e.g., demographics, psychiatric disorders) and methodologically (e.g., MRI protocols, software, statistical analyses). Despite the high network replicability observed across studies, this heterogeneity may have led to an underestimation of the true level of replicability. In the future, researchers may wish to explore the variability of the identified networks across more homogeneous subgroups. Second, we used normative resting-state functional connectivity and structural covariance data from a large sample of healthy individuals (n = 1,000, 50% female) to map heterogeneous brain regions onto a common network. Consequently, it is possible that differences in demographics between the included case samples and the normative sample may have resulted in some variations. Third, we relied on a meta-analytic rather than a mega-analytic approach. Again, international large-scale collaborative initiatives such as the ENIGMA-Antisocial Behavior Working Group ^112^, where individual-level data are shared, may provide a better understanding of the neurobiological correlates of aggression compared to meta-analytic approaches. Despite these limitations, our work offers a promising framework to overcome previous concerns about replicability in psychiatric neuroscience, while also providing a bridge for integrating multiple levels of analysis.

## Conclusion

In this work, we empirically delineated functional and structural networks associated with aggression using normative network mapping. While functional studies were dominated by a subcortical network, structural ones showed the highest overlap in areas of the ventral attention network such as the amygdala and insula. Yet, both functional and structural studies converged in areas of the default mode network such as the medial prefrontal cortex. Further, we identified a spatially generalizable transcriptomic signature of aggression, wherein genes involved in synaptic function and genomic regulation contributed most to the covariance between gene expression maps and aggression-related brain networks. Future studies may further refine these models by integrating single-cell transcriptomic data, developmental trajectories, and gene-environment interactions to delineate the causal pathways linking the multiple levels of analysis that underlie aggressive behavior.

## Data Availability

Peak coordinates of neuroimaging studies included in the meta-analysis will be made available upon reasonable request by the corresponding author. Preprocessed resting-state data from the Brain Genomics Superstruct Project is available on the Harvard Dataverse (https://www.neuroinfo.org/gsp). Voxelwise composite score maps (gene expression and imaging modalities) were generated by projecting PLS-derived weights into voxel space. These maps, along with unthresholded aggression brain network maps, will be made publicly available as a collection on NeuroVault upon publication.

## Code Availability

Coordinate-based meta-analysis ALE was conducted using GingerALE version 3.0.2. (http://www.brainmap.org/ale). Network Mapping was conducted using scripts from Stubbs and colleagues ^34^ (https://github.com/nimlab/NMH_Stubbs2023). Gene expression patterns were obtained from the Allen Human Brain Atlas ^45, 65^ (https://github.com/rmarkello/abagen).

## Acknowledgements

This study did not receive any specific funding. SADB was supported by an Economic and Social Research Council Grant (ES/V003526/1), and MBR was supported by a Brain and Behavior Research Foundation Young Investigator Award (30919).

