## Supplementary Material for "The neural orchestra of aggression: neurogenetic network mapping of human aggressiveness"

– SYSTEMATIC REVIEW AND META-ANALYSIS –

Jules Roger Dugré, PhD

Department of Psychology, College of Literature, Science, and the Arts, University of Michigan, Ann Arbor, MI, USA 48109

&

Stéphane Alexandre De Brito, PhD;

Centre for Human Brain Health, University of Birmingham, School of Psychology, Birmingham, UK B15 2TT

### **Supplementary Method****s**

#### MRI data acquisition – GSP1000 dataset

The GSP1000 cohort contains a fully 1:1 M:F matched dataset of one thousand human participants ages 18 to 35 years that were scanned with a Siemens Tim Trio 3T scanners (Siemens Healthcare, Erlangen, Germany) at Harvard University and Massachusetts General Hospital using the vendor-supplied 12-channel phased-array head coil. Participants were matched based on age and sex. High-resolution (1.2 mm isotropic) multi-echo MPRAGE sequences were acquired for a duration of 2 min 12s, with sagittal acquisition orientation, 144 slices, TR=2200 ms, TE=1.5/3.4/5.2/7.0 ms, FA= 7°, TI=1100 ms, Resolution =1.2×1.2×1.2 mm.

For functional data, subjects were instructed to remain still, stay awake, and keep their eyes open. EPI parameters were as follows: repetition time (TR) = 3,000 ms, echo time (TE) = 30 ms, flip angle (FA) = 85°, 3 × 3 × 3-mm voxels, field of view (FOV) = 216, and 47 axial slices collected with interleaved acquisition and no gap between slices. Each functional run lasted 6.2 min (124 time points). One or two runs were acquired per subject (average of 1.7 runs).

### **Supplementary Tables**

#### **Supplementary Table 1.** PRISMA 2020 Checklist

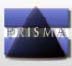
**PRISMA 2020 Checklist**

| **Section and Topic** | **Item #** | **Checklist item** | **Location where item is reported** |
| --- | --- | --- | --- |
| **TITLE** | | |  |
| Title | 1 | Identify the report as a systematic review. | No |
| **ABSTRACT** | | |  |
| Abstract | 2 | See the PRISMA 2020 for Abstracts checklist. | S Table2 |
| **INTRODUCTION** | | |  |
| Rationale | 3 | Describe the rationale for the review in the context of existing knowledge. | 3-6 |
| Objectives | 4 | Provide an explicit statement of the objective(s) or question(s) the review addresses. | 6 |
| **METHODS** | | |  |
| Eligibility criteria | 5 | Specify the inclusion and exclusion criteria for the review and how studies were grouped for the syntheses. | 24 |
| Information sources | 6 | Specify all databases, registers, websites, organisations, reference lists and other sources searched or consulted to identify studies. Specify the date when each source was last searched or consulted. | 23 |
| Search strategy | 7 | Present the full search strategies for all databases, registers and websites, including any filters and limits used. | 23 |
| Selection process | 8 | Specify the methods used to decide whether a study met the inclusion criteria of the review, including how many reviewers screened each record and each report retrieved, whether they worked independently, and if applicable, details of automation tools used in the process. | 24 |
| Data collection process | 9 | Specify the methods used to collect data from reports, including how many reviewers collected data from each report, whether they worked independently, any processes for obtaining or confirming data from study investigators, and if applicable, details of automation tools used in the process. | NA |
| Data items | 10a | List and define all outcomes for which data were sought. Specify whether all results that were compatible with each outcome domain in each study were sought (e.g. for all measures, time points, analyses), and if not, the methods used to decide which results to collect. | 24 |
|  | 10b | List and define all other variables for which data were sought (e.g. participant and intervention characteristics, funding sources). Describe any assumptions made about any missing or unclear information. | NA |
| Study risk of bias assessment | 11 | Specify the methods used to assess risk of bias in the included studies, including details of the tool(s) used, how many reviewers assessed each study and whether they worked independently, and if applicable, details of automation tools used in the process. | NA |
| Effect measures | 12 | Specify for each outcome the effect measure(s) (e.g. risk ratio, mean difference) used in the synthesis or presentation of results. | NA |
| Synthesis methods | 13a | Describe the processes used to decide which studies were eligible for each synthesis (e.g. tabulating the study intervention characteristics and comparing against the planned groups for each synthesis (item #5)). | NA |
|  | 13b | Describe any methods required to prepare the data for presentation or synthesis, such as handling of missing summary statistics, or data conversions. | NA |
|  | 13c | Describe any methods used to tabulate or visually display results of individual studies and syntheses. | NA |
|  | 13d | Describe any methods used to synthesize results and provide a rationale for the choice(s). If meta-analysis was performed, describe the model(s), method(s) to identify the presence and extent of statistical heterogeneity, and software package(s) used. | NA |
|  | 13e | Describe any methods used to explore possible causes of heterogeneity among study results (e.g. subgroup analysis, meta-regression). | NA |
|  | 13f | Describe any sensitivity analyses conducted to assess robustness of the synthesized results. | NA |
| Reporting bias assessment | 14 | Describe any methods used to assess risk of bias due to missing results in a synthesis (arising from reporting biases). | NA |
| Certainty assessment | 15 | Describe any methods used to assess certainty (or confidence) in the body of evidence for an outcome. | 26 |
| **RESULTS** | | |  |
| Study selection | 16a | Describe the results of the search and selection process, from the number of records identified in the search to the number of studies included in the review, ideally using a flow diagram. | Fig.1 |
|  | 16b | Cite studies that might appear to meet the inclusion criteria, but which were excluded, and explain why they were excluded. | NA |
| Study characteristics | 17 | Cite each included study and present its characteristics. | Stable3  STable4 |
| Risk of bias in studies | 18 | Present assessments of risk of bias for each included study. | NA |
| Results of individual studies | 19 | For all outcomes, present, for each study: (a) summary statistics for each group (where appropriate) and (b) an effect estimate and its precision (e.g. confidence/credible interval), ideally using structured tables or plots. | NA |
| Results of syntheses | 20a | For each synthesis, briefly summarise the characteristics and risk of bias among contributing studies. | NA |
|  | 20b | Present results of all statistical syntheses conducted. If meta-analysis was done, present for each the summary estimate and its precision (e.g. confidence/credible interval) and measures of statistical heterogeneity. If comparing groups, describe the direction of the effect. | 8-12 |
|  | 20c | Present results of all investigations of possible causes of heterogeneity among study results. | NA |
|  | 20d | Present results of all sensitivity analyses conducted to assess the robustness of the synthesized results. | 8-12 |
| Reporting biases | 21 | Present assessments of risk of bias due to missing results (arising from reporting biases) for each synthesis assessed. | NA |
| Certainty of evidence | 22 | Present assessments of certainty (or confidence) in the body of evidence for each outcome assessed. | NA |
| **DISCUSSION** | | |  |
| Discussion | 23a | Provide a general interpretation of the results in the context of other evidence. | 19-23 |
|  | 23b | Discuss any limitations of the evidence included in the review. | 22-23 |
|  | 23c | Discuss any limitations of the review processes used. | 22-23 |
|  | 23d | Discuss implications of the results for practice, policy, and future research. | 21 |
| **OTHER INFORMATION** | | |  |
| Registration and protocol | 24a | Provide registration information for the review, including register name and registration number, or state that the review was not registered. | NA |
|  | 24b | Indicate where the review protocol can be accessed, or state that a protocol was not prepared. | NA |
|  | 24c | Describe and explain any amendments to information provided at registration or in the protocol. | NA |
| Support | 25 | Describe sources of financial or non-financial support for the review, and the role of the funders or sponsors in the review. | 31 |
| Competing interests | 26 | Declare any competing interests of review authors. | 31 |
| Availability of data, code and other materials | 27 | Report which of the following are publicly available and where they can be found: template data collection forms; data extracted from included studies; data used for all analyses; analytic code; any other materials used in the review. | 30-31 |

*From:* Page MJ, McKenzie JE, Bossuyt PM, Boutron I, Hoffmann TC, Mulrow CD, et al. The PRISMA 2020 statement: an updated guideline for reporting systematic reviews. BMJ 2021;372:n71. doi: 10.1136/bmj.n71. This work is licensed under CC BY 4.0. To view a copy of this license, visit <https://creativecommons.org/licenses/by/4.0/>

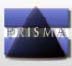
**PRISMA 2020 for Abstracts Checklist**

#### **Supplementary Table 2.** PRISMA 2020 Abstracts Checklist

| **Section and Topic** | **Item #** | **Checklist item** | **Reported (Yes/No)** |
| --- | --- | --- | --- |
| **TITLE** | | |  |
| Title | 1 | Identify the report as a systematic review. | Yes |
| **BACKGROUND** | | |  |
| Objectives | 2 | Provide an explicit statement of the main objective(s) or question(s) the review addresses. | Yes |
| **METHODS** | | |  |
| Eligibility criteria | 3 | Specify the inclusion and exclusion criteria for the review. | No |
| Information sources | 4 | Specify the information sources (e.g. databases, registers) used to identify studies and the date when each was last searched. | No |
| Risk of bias | 5 | Specify the methods used to assess risk of bias in the included studies. | No |
| Synthesis of results | 6 | Specify the methods used to present and synthesise results. | No |
| **RESULTS** | | |  |
| Included studies | 7 | Give the total number of included studies and participants and summarise relevant characteristics of studies. | Yes |
| Synthesis of results | 8 | Present results for main outcomes, preferably indicating the number of included studies and participants for each. If meta-analysis was done, report the summary estimate and confidence/credible interval. If comparing groups, indicate the direction of the effect (i.e. which group is favoured). | Yes |
| **DISCUSSION** | | |  |
| Limitations of evidence | 9 | Provide a brief summary of the limitations of the evidence included in the review (e.g. study risk of bias, inconsistency and imprecision). | No |
| Interpretation | 10 | Provide a general interpretation of the results and important implications. | Yes |
| **OTHER** | | |  |
| Funding | 11 | Specify the primary source of funding for the review. | NA |
| Registration | 12 | Provide the register name and registration number. | NA |

| **Supplementary Table 3.** Description of the Included Voxel-Based Morphometry Studies on Trait Aggression | | | | | | | | |
| --- | --- | --- | --- | --- | --- | --- | --- | --- |
| First Author, Date |  | Sample of Interest | | | | Controls (n=) | Analysis | Aggression Measures |
|  | Location | Group Description | n | Mean Age | Males (%) |  |  |  |
| (Bertsch *et al.*, 2013) | Germany | Offenders with ASPD + BPD | 13 | 28.9 | 100.0% | 14 | Group | FAF-Total;  FAF-Spontaneous;  FAF-Reactive |
|  |  | Offenders with ASPD + Psychopathic Traits | 12 | 27.3 | 100.0% | - |  |  |
| (Besteher *et al.*, 2017) | Germany, Italy | Population-Based | 349 | 30.6 | 52.7% | - | Dimension | SCL90-R Aggression |
| (Bobes *et al.*, 2013) | Mexico | Violent Group (Community) | 25 | 30.6 | 100.0% | 29 | Group | RPQ-Total;  RPQ-Reactive;  RPQ-Proactive;  BDHI Total |
| (Breitschuh *et al.*, 2018) | Germany | Martial Artists | 21 | 26.1 | 100.0% | 26 | Group | FAF-Aggression;  FAF-Spontaneous;  FAF-Reactive |
| (Budhiraja *et al.*, 2017) | Sweden | Females with Conduct Disorder | 31 | 24.1 | 0.0% | 25 | Group | Aggressive CD Count |
| (Chester *et al.*, 2017) | United States | Undergrads/Community | 138 | 19.4 | 34.1% | - | Dimension | BAQ-Physical;  BAQ-Verbal |
| (Coccaro *et al.*, 2016) | United States | Intermittent Explosive Disorder | 57 | 34.4 | 54.0% | 111 | Group | LHA-Aggression;  BPAQ |
| (Coccaro *et al.*, 2018) | United States | Healthy Subjects | 287 | 35.0 | 50.5% | - | Dimension | LHA-Aggression |
| (Dalwani *et al.*, 2011) | United States | Adolescents with Substance and Conduct Problems | 25 | 16.6 | 100.0% | 19 | Group | Peak Aggression Scale |
| (Dalwani *et al.*, 2015) | United States | Adolescents with severe CD | 22 | 16.1 | 0.0% | 21 | Group | Peak Aggression Scale |
| (Drachman *et al.*, 2022) | United States | Bipolar Disorder | 38 | 35.7 | 26.0% | - | Dimension | BGA |
| (Fairchild *et al.*, 2011) | United Kingdom | Early-Onset Conduct Disorder | 36 | 17.7 | 100.0% | 27 | Group | Aggressive CD Count |
|  |  | Adolescent-Onset Conduct Disorder | 27 | 17.9 | 100.0% | - |  |  |
| (Fairchild *et al.*, 2013) | United Kingdom | Females with Conduct Disorder | 22 | 17.2 | 0.0% | 20 | Group | Aggressive CD Count |
|  |  | Adolescents with CD | 44 | 17.4 | 50.0% | 40 |  |  |
| (Gao *et al.*, 2022) | China | TD with Childhood Maltreatment | 46 | 15.2 | 100.0% | 40 | Group | BPAQ-Total;  BPAQ-Physical;  BPAQ-Verbal |
|  |  | CD with Childhood Maltreatment | 62 | 14.8 | 100.0% | 34 |  |  |
| (Gong *et al.*, 2023) | China | Undergraduates | 176 | 20.6 | 47.2% | - | Dimension | DD - Machiavellianism |
| (Gregory *et al.*, 2012) | United Kingdom | ASPD+Psychopathy | 17 | 38.9 | 100.0% | 22 | Group | RPQ-Total;  RPQ-Proactive;  RPQ-Reactive |
|  |  | ASPD-Psychopathy | 27 | 36.1 | 100.0% | - |  |  |
| (Hofhansel *et al.*, 2020) | Germany | Offenders | 27 | 35.6 | 100.0% | - | Dimension | RPQ-Total;  RPQ-Reactive;  RPQ-Proactive;  BPAQ-Total;  BPAQ-Physical;  BPAQ-Verbal |
|  |  | Healthy Controls | 27 | 34.4 | 100.0% | - |  |  |
| (Huber *et al.*, 2018) | Switzerland | Agitated-Aggressive Group (ARMS+FEP) | 49 | 25.5 | 75.5% | 87 | Group | BPRS-EC |
| (Ibrahim *et al.*, 2021) | United States | Children with DBDs | 88 | 11.7 | 65.9% | 50 | Group | CBCL-AGG |
| (Kuhlmann *et al.*, 2013) | Germany | Women with BPD | 30 | 23.7 | 0.0% | 33 | Group | FAF-Aggression;  FAF-Spontaneous;  FAF-Reactive;  STAXI-Anger OUT |
| (Liu *et al.*, 2020) | China | Violent Patients with Schizophrenia | 48 | 29.0 | 100.0% | 55 | Group | MOAS |
| (Lui *et al.*, 2009) | China | Schizophrenia Patients | 68 | 24.2 | 44.1% | 68 | Group | PANSS - IMP/AGG |
| (Martinez-Horta *et al.*, 2021) | Spain | Irritable/Aggressive Group (Huntington Disease) | 14 | 53.7 | 28.6% | 17 | Group | PBA-s - Aggression |
| (Mohammadi *et al.*, 2020) | Germany | Gamers (Violent Video Game) | 29 | 23.6 | 100.0% | 29 | Both | FAF-Reactive |
| (Raschle *et al.*, 2018) | Multi-Center (Europe) | Typically Developing Girls | 108 | 13.9 | 0.0% | 81 | Group | CBCL-AGG |
| (Romero-Rebollar *et al.*, 2015) | Mexico | Violent Subjects (Community) | 23 | 29.4 | 100.0% | 24 | Group | RPQ-Reactive;  BDHI Total |
| (Seok and Cheong, 2020) | Korea | Intermittent Explosive Disorder | 15 | 28.5 | 100.0% | 15 | Both | LHA-Aggression;  BPAQ |
| (Schiffer *et al.*, 2011) | Germany | Individuals with SUDs | 25 | 36.9 | 100.0% | 26 | Group | LHA |
| (Schiffer *et al.*, 2013) | Germany | Schizophrenia Patients with CD | 27 | 36.2 | 100.0% | 23 | Group | LHA-AGG |
|  |  | Adults with CD | 27 | 36.0 | 100.0% | 25 |  |  |
| (Schoretsanitis *et al.*, 2019) | Switzerland | Schizophrenia Patients | 84 | 36.7 | 64.3% | - | Dimension | MOAS-Total;  MOAS-Verbal;  MOAS-Physical |
| (Soloff *et al.*, 2008) | United States | BPD Females | 22 | 26.1 | 0.0% | 19 | Group | LHA-AGG |
|  |  | BPD Males | 12 | 30.0 | 100.0% | 11 |  |  |
| (Soloff *et al.*, 2014) | United States | BPD with High Lethality of Suicidal Attempt | 16 | 36.1 | 68.8% | - | Dimension | LHA-Aggression |
|  |  | BPD with Low Lethality of Suicidal Attempt | 35 | 27.4 | 85.7% | - |  |  |
| (Sterzer *et al.*, 2007) | Germany | Conduct Disorder | 12 | 12.8 | 100.0% | 12 | Group | CBCL-AGG (T-score) |
| (Wang *et al.*, 2023) | China | Undergraduates | 202 | 20.4 | 37.6% | - | Dimension | BPAQ-Total;  BPAQ-Physical;  BPAQ-Verbal |
| (Woermann *et al.*, 2000) | United Kingdom | TLE with IED | 24 | 27.0 | 68.0% | 59 | Group | SDAS-9 |
| (Yang *et al.*, 2017) | United States | Adolescents Twins (Girls) | 52 | 14.0 | 0.0% | - | Dimension | RPQ-Total;  RPQ-Reactive;  RPQ-Proactive |
|  |  | Adolescents Twins (Boys) | 54 | 14.0 | 100.0% | - |  |  |
| Note. ARMS = At Risk Mental State; FEP = First Episode Psychosis; TLE = Temporal Lobe Epilepsy; CD = Conduct Disorder; SUD = Substance Use Disorder; BPD = Borderline Personality Disorder; DBD = Disruptive Behaviors Disorders; TD = Typically Developping. Aggressive CD count = Diagnostic Criteria (APA, 2013); BAQ = Brief Aggression Questionnaire (Webster *et al.*, 2015); BDHI = Buss-Durkee Hostility Inventory (Buss and Durkee, 1957); BGA = Brown Goodwin Aggression Questionnaire (Brown *et al.*, 1979); BPAQ = Buss-Perry Aggression Questionnaire (Buss and Perry, 1992); BPRS-Excitement = Brief Psychiatric Rating Scale (Overall and Gorham, 1988); CBCL-AGG = Child Behavior Checklist – Aggression Syndrome Scale (Achenbach and Rescorla, 2000); DD-Machiavellianism = Dirty Dozen (Jonason and Webster, 2010); FAF/FAI = Factors of Aggression Questionnaire (Hampel and Selg, 1975); LHA = Life History of Aggression (Coccaro *et al.*, 1997); MOAS = Modified Overt Aggression Scale (Sorgi *et al.*, 1991); PANSS = Positive and Negative Syndrome Scale for Schizophrenia (Kay *et al.*, 1987); PABRS = Peak Aggressive Behavior Rating Scale (Crowley *et al.*, 2001); PBA-s = Problem Behaviors Assessment for Huntington Disease (Craufurd *et al.*, 2001); RPQ = Reactive & Proactive Aggression(Dodge and Coie, 1987, Raine *et al.*, 2006); SCL-90 = The Symptom Checklist-90 (Derogatis *et al.*, 1973); SDAS-9 = Social Dysfunction and Aggression Scale (Wistedt *et al.*, 1990); STAXI-AX-OUT = State-Trait Anger Expression Inventory - Anger Expression OUT (Spielberger *et al.*, 1999); | | | | | | | | |

| **Supplementary Table 4.** Description of the Included Task-Based Functional MRI Studies on Trait Aggression | | | | | | | | |
| --- | --- | --- | --- | --- | --- | --- | --- | --- |
| First Author, Date | Sample of Interest | | | | Controls (n=) | Analysis | Aggression Measures | fMRI Task |
|  | Group Description | n | Mean Age | Males (%) |  |  |  |  |
| Aggensteiner, 2020^60^ | Oppositional Defiant Disorder/Conduct Disorder | 108 | 13.2 | 82.4% | 69 | Case Control | CBCL-AGG;  RPQ-Total;  RPQ-Reactive;  RPQ-Proactive | Emotional Faces |
| Aghajani, 2021^61^ | Conduct Disorder with Limited Prosocial Emotions | 19 | 16.4 | 100.0% | 31 | Case Control | RPQ-Total;  RPQ-Proactive;  RPQ-Reactive | Emotional Recognition & Emotional Resonance |
| Beames, 2020^62^ | Healthy Subjects | 21 | 21.0 | 57.1% | 24 | Both | BPAQ-Total | Unsolvable Anagram |
| Bertsch, 2018^63^ | Borderline Personality Disorder | 30 | 26.9 | 0.0% | 28 | Case Control | STAXI AX-OUT | Approach Avoidance Task (Faces) |
| Bertsch, 2022^64^ | Borderline Personality Disorder | 48 | 29.6 | 0.0% | 28 | Case-Control | BPAQ-Total | Social Threat Aggression  Paradigm |
| Blair, 2021^65^ | Adolescents from Residential Care Facility | 98 | 16.0 | 69.4% | - | Dimension | Aggression Incident Reports | The Looming Task |
| Bobes, 2013^66^ | Violent Men in Community | 25 | 30.6 | 100.0% | 29 | Case Control | RPQ-Total;  RPQ-Proactive;  RPQ-Reactive;  BDHI-Total | Fearful Faces |
| Bubenzer-Busch,  2016^67^ | Attention-Deficit/Hyperactivity Disorder & Disruptive Behavior Disorders | 27 | 10.9 | 100.0% | 27 | Both | CBCL-AGG;  BPAQ-Total;  BPAQ-Physical;  BPAQ-Verbal;  RPQ-Total;  RPQ-Reactive;  RPQ-Proactive | mPSAG |
| Coccarro, 2007^68^ | Intermittent Explosive Disorder | 10 | 34.3 | 50.0% | 10 | Case-Control | LHA-AGG;  BPAQ-Total | Emotional Faces |
| Coccaro, 2022^69^ | Intermittent Explosive Disorder | 19 | 35.0 | 42.0% | 26 | Case Control | LHA-AGG;  BPAQ-AGG | V-SEIP task |
| Coccaro, 2021^70^ | Healthy Subjects | 26 | 32.0 | 50.0% | - | Dimension | LHA-AGG | V-SEIP task |
| Cohn, 2013^71^ | Disruptive Behavior Disorders (Desisters) | 25 | 17.6 | 80.0% | 26 | Case Control | RPQ-Total | Fear Conditioning Task |
|  | Disruptive Behavior Disorders (Persisters) | 25 | 17.3 | 72.0% | - |  |  |  |
| Crowley, 2010^72^ | Adolescents with Antisocial Substance Disorder | 20 | 16.5 | 100.0% | 20 | Case-Control | Peak Aggressive Behavior Rating Scale | Colorado Balloon Game |
| Crowley, 2015^73^ | Males with Antisocial Substance Disorder | 20 | 16.5 | 100.0% | 20 | Case-Control | Peak Aggressive Behavior Rating Scale | Decision-Making Behavioral Task |
|  | Females with Antisocial Substance Disorder | 21 | 16.2 | 0.0% | 20 |  |  |  |
| Cremers, 2016^74^ | Intermittent Explosive Disorder | 17 | 32.5 | 58.8% | 14 | Case Control | LHA-AGG | Emotional Faces |
| Crum, 2023^75^ | Adolescents with Substance Misuse & Rule-Breaking Problems | 55 | 16.7 | 60.0% | 125 | Case Control | CBCL-AGG | Passive Avoidance Task |
| Decety, 2009^76^ | Conduct Disorder | 8 | 16 to 18 | NA | 8 | Case-Control | Aggressive CD Symptom Count | Empathy for Pain |
| Eijsker, 2019^77^ | Patients with Misophonia | 22 | 33.2 | 27.0% | 21 | Case-Control | BPAQ-Total;  BPAQ-Physical; BPAQ-Verbal | Stop Signal Task |
| Fairchild, 2014^78^ | Conduct Disorder | 20 | 17.0 | 0.0% | 20 | Case Control | Aggressive CD Symptom Count | Emotional Faces |
| Gan, 2016^79^ | Clinical & Sub-Clinical Intermittent Explosive Disorder | 9 | 34.4 | 100.0% | 9 | Case Control | STAXI2-AX-OUT | PSAP |
| García-Martí 2013^80^ | Treatment Resistant Schizophrenia with Auditory Hallucination | 32 | NA | NA | - | Dimension | BPRS-Hostility | Emotional Words (Auditory) |
| Gatze-Kopp, 2009^81^ | Externalizing Disorders | 19 | 13.6 | 100.0% | 11 | Case-Control | CBCL-AGG | MIDT |
| Gregory, 2015^82^ | Antisocial Personality Disorder (without Psychopathy) | 20 | 36.8 | 100.0% | 18 | Case Control | RPQ-Total;  RPQ-Reactive;  RPQ-Proactive | Probabilistic Response Reversal Task |
|  | Antisocial Personality Disorder (with Psychopathy) | 12 | 40.1 | 100.0% | - |  |  |  |
| Hazlett, 2012^83^ | Borderline Personality Disorder | 33 | 31.6 | 39.0% | 32 | Case-Control | BPAQ-Total | Repeated Emotional Pictures |
|  | Schizotypal Personality Disorder | 28 | 35.9 | 57.0% | - |  |  |  |
| Heesink, 2017^84^ | Veterans with Anger Problems | 27 | 36.4 | 100.0% | 30 | Case Control | BPAQ-Physical;  BPAQ-Verbal | Fear-and-Escape Task (FAET) |
| Heesink, 2018^85^ | Veterans with Anger Problems | **30** | 36.3 | 100.0% | 29 | Case Control | BPAQ-Physical;  BPAQ-Verbal | Affective Stimuli (IAPS) |
| Herpertz, 2008^86^ | Childhood-Onset Conduct Disorder | 22 | 14.7 | 100.0% | 22 | Case Control | CBCL-AGG | Affective Stimuli (IAPS) |
|  | Attention-Deficit/Hyperactivity Disorder | 13 | 14.0 | 100.0% | 13 |  |  |  |
| Herpertz, 2017^87^ | Borderline Personality Disorder | 33 | 26.2 | 0.0% | 30 | Case-Control | BPAQ-Total | Anger-Aggression Scripts |
|  | Borderline Personality Disorder | 23 | 30.7 | 100.0% | 26 |  |  |  |
| Ibrahim, 2019^88^ | Autism Spectrum Disorder/Disruptive Behavior Disorders | 18 | 12.7 | 88.9% | 19 | Case-Control | CBCL-AGG | Emotional Faces |
| Jakobi, 2022^89^ | Attention-Deficit/Hyperactivity Disorder | 78 | 34.2 | 43.6% | 78 | Both | RPQ-Proactive;  RPQ-Reactive | Emotional Faces |
| Jiang, 2018^90^ | Healthy Controls | 19 | 20.0 | 52.6% | 20 | Case Control | BPAQ-Total | TAP |
| Kesner, 2020^91^ | Medical Students (High Xenophobic) | 19 | 23.0 | 52.6% | 19 | Case Control | BPAQ-Total;  BPAQ-Physical; BPAQ-Verbal | Affective Stimuli (Refugees/Terrorists) |
| Kim, 2018^92^ | Delinquent Adolescents | 8 | 14.5 | 75.0% | 17 | Case-Control | CBCL-AGG | Rest |
| Konzok, 2021^93^ | Healthy Subjects (High Externalizing Traits) | 31 | 23.1 | 50.8% | 30 | Case Control | TriPM-Meanness | ScanSTRESS |
| Konzok, 2022^94^ | Healthy Subjects (High Externalizing Traits) | 32 | 23.6 | 50.0% | 31 | Case Control | TriPM-Meanness,  K-FAF-Spontaneous; K-FAF-Reactive;  BPAQ-Physical;  BPAQ-Verbal;  RPQ-Proactive;  RPQ-Reactive | mTAP |
| Krauch, 2018^95^ | Borderline Personality Disorder | 20 | 16.4 | 0.0% | 20 | Case Control | BPAQ-Total | Anger-Aggression Scripts |
|  | Borderline Personality Disorder | 34 | 25.7 | 0.0% | 32 |  |  |  |
| Kumari, 2006^96^ | Violent Patients with Schizophrenia | 12 | 34.0 | 100.0% | 13 | Case-Control | Gunn–Robertson Scale | N-Back Task |
|  | Antisocial Personality Disorder | 10 | 31.3 | 100.0% | 13 |  |  |  |
| Kumari, 2009^97^ | Violent Patients with Schizophrenia | 13 | 34.5 | 100.0% | 13 | Case Control | Gunn–Robertson Scale | Fear Elicitation (Shock) |
|  | Antisocial Personality Disorder | 13 | 32.9 | 100.0% | 14 |  |  |  |
| Li,2020^98^ | Primary School Students | 77 | 10.2 | 45.5% | - | Dimension | BWAQ-Total;  BWAQ-Physical;  BWAQ-Verbal | Rest |
| Martinelli, 2021^99^ | Healthy Subjects | 50 | 15.6 | 50.0% | - | Dimension | BPAQ-Physical | Emotional Faces |
| Mathur, 2023^100^ | Residential Treatment Program | 42 | 16.2 | 61.0% | 41 | Case-Control | RPQ-Total^9^;  RPQ-Reactive^9^;  RPQ-Proactive^9^ | Retaliation Task |
| McCloskey, 2016^101^ | Intermittent Explosive Disorder | 20 | 33.2 | 60.0% | 20 | Case-Control | LHA-AGG | Emotional Faces |
| Michalska 2016^102^ | Child from Mental Health and Pediatric Clinics | 107 | 10.1 | 48.0% | - | Dimension | RPQ-Reactive | Harm scenarios |
| Moeller, 2014^103^ | Intermittent Explosive Disorder | 11 | 33.5 | 100.0% | 38 | Case-Control | STAXI AX-OUT | Stroop Task |
| Murray, 2023^104^ | Youths from Low Income Families | 128 | 15.9 | 42.0% | - | Dimension | CBCL-AGG | MIDT |
| Passamonti, 2010^105^ | Early-Onset Conduct Disorder | 11 | 17.7 | 100.0% | 18 | Case Control | Aggressive CD Symptom Count | Emotional Faces |
|  | Adolescence-Onset Conduct Disorder | 11 | 17.1 | 100.0% | - |  |  |  |
| Pawliczek, 2013^106^ | Students (High Trait Aggression) | 21 | 22.2 | 100.0% | 18 | Case-Control | BPAQ-Total;  BPAQ-Physical;  BPAQ-Verbal;  FAI-Total;  FAI-Spontaneous;  FAI-Reactive | Unsolvable Anagrams |
| Pawliczek, 2013^107^ | University Students (High Externalizing Traits) | 17 | 22.2 | 100.0% | 16 | Case Control | BPAQ-Total;  BPAQ-Physical;  BPAQ-Verbal | Stop Signal (Affective Faces) |
| Perino, 2019^108^ | Adolescents (with School or Legal intervention) | 24 | 16.2 | 50.0% | - | Dimension | University of Illinois Bully Scale | The Catch Game |
| Prehn, 2013^109^ | Borderline Personality Disorder/Antisocial Personality Disorder | 15 | 27.9 | 100.0% | 17 | Case Control | FAF-Aggression;  FAF-Spontaneous;  FAF-Reactive | N-Back (w/wo affective stimuli) |
| Prehn, 2013^110^ | Emotionally Hyporeactive Offenders | 11 | 27.6 | 100.0% | 13 | Case-Control | FAF-Spontaneous; FAF-Reactive | Monetary Decision-Making Task |
|  | Emotionally Hyperreactive Offenders | 12 | 27.8 | 100.0% | - |  |  |  |
| Rahrig, 2021^111^ | Active Coping Group | 11 | 35.1 | 45.0% | 9 | Case Control | BAQ | TAP |
| Repple, 2018^112^ | Healthy Controls | 22 | 24.8 | 100.0% | 20 | Case Control | BPAQ-Total;  BPAQ-Physical;  BPAQ-Verbal | mTAP |
| Schröder, 2019^113^ | Patients with Misophonia | 21 | 33.1 | 28.0% | 23 | Case-Control | BPAQ-Total;  BPAQ-Physical; BPAQ-Verbal | Clip Viewing |
| Seok, 2020^114^ | Intermittent Explosive Disorder | 15 | 28.5 | 100.0% | 15 | Both | LHA-AGG;  BPAQ-Total | Affective Videos |
| Sethi, 2018^115^ | Primary Psychopathy (Community) | 50 | 19.8 | 82.0% | 82 | Case-Control | BPAQ-Total | Match-To-Sample (Faces) |
|  | Secondary Psychopathy (Community) | 100 | 19.5 | 58.0% | - |  |  |  |
| Shao, 2017^116^ | University Students (High Psychopathic Traits) | 29 | 20.4 | 48.3% | 23 | Case-Control | PPI-SCI | Face Familiarity |
| Soloff, 2017^117^ | Borderline Personality Disorder | 31 | 30.0 | 0.0% | - | Dimension | LHA-AGG | Go/No-Go Task |
| Szycik, 2017^118^ | Violent Video Game Users | 15 | 22.8 | 100.0% | 15 | Case Control | K-FAF-AGG | Socio-Affective Situations |
| Taubner, 2021^119^ | Violent Offenders | 25 | 19.9 | 100.0% | - | DImension | RPQ-Total | Interactive Video |
|  | Healthy Controls | 21 | 20.0 | 100.0% | - |  |  |  |
| Tonnaer, 2017^120^ | Violent Offenders | 16 | 35.8 | 100.0% | 18 | Case-Control | RPQ-Total;  RPQ-Reactive;  RPQ-Proactive;  BPAQ-Total | Emotional Stories |
| Vanova, 2022^121^ | Healthy Subjects (University Network) | 22 | 24.1 | 31.8% | - | Dimension | TriPM-Meanness | Lexical Decision Task |
| Weidler, 2019^122^ | OPRM genotype (G-) | 39 | 25.2 | 100.0% | 20 | Case Control | BPAQ-Total;  BPAQ-Physical;  BPAQ-Verbal;  RPQ-Proactive;  RPQ-Reactive | TAP |
| White, 2016^123^ | Disruptive Behavior Disorders | 30 | 15.0 | 63.3% | 26 | Case Control | RPQ-Proactive^9^;  RPQ-Reactive^9^ | Social Fairness Game |
| White, 2018^124^ | Disruptive Behavior Disorders | 31 | 14.6 | 71.0% | 27 | Case-Control | RPQ-Reactive^9^ | The Looming Task |
| Yoder, 2015^125^ | Healthy Subjects who Watch Mixed Martial Arts | 43 | 25.0 | 100.0% | - | Dimension | PPI-SCI | Passive Viewing MMA |
| Zhang, 2021^126^ | Conduct Disorder | 101 | 15.9 | 64.4% | 77 | Case-Control | RPQ-Total;  RPQ-Reactive;  RPQ-Proactive | Passive Avoidance Task |
| Note. CBCL-AGG = Child Behavior Checklist – Aggression Syndrome Scale ^127^; RPQ = Reactive & Proactive Aggression^9, 12^; BPAQ = Buss-Perry Aggression Questionnaire ^16^; STAXI-AX-OUT = State-Trait Anger Expression Inventory - Anger Expression OUT ^128^; LHA = Life History of Aggression ^129^; BWAQ = Buss-Warren Aggression Questionaire ^130^; TriPM = Triarchic Psychopathy Measure ^131^; K-FAF = Short version of the Factors of Aggression Questionnaire ^132^; BDHI = Buss-Durkee Hostility Inventory ^15^; BAQ = Brief Aggression Questionnaire ^133^; FAF/FAI = Factors of Aggression Questionnaire ^134^; Peak Aggressive Behavior Rating Scale ^135^; BPRS-Hostility = Brief Psychiatric Rating Scale ^136^; Gunn-Robertson Violence Scale ^137^; University of Illinois Bully Scale ^138^; PPI-SCI = Psychopathic Personality Inventory-Self-Centered Impulsivity ^139^ | | | | | | | | |

| **Supplementary Table 5.** Most replicable peaks of the Task-based Functional Brain Network underpinning Trait Aggression | | | | | |
| --- | --- | --- | --- | --- | --- |
| Regions | MNI Coordinates | | | Peak Intensity (t-values) | Replicability (N, %) |
|  | x | y | z |  |  |
| Caudate Nucleus | -8 | 6 | 6 | 3.64 | 33 (84.62 %) |
| Caudate Nucleus | 10 | 10 | 4 | 5.22 | 30 (76.92 %) |
| perigenual part of the Anterior Cingulate Cortex | -4 | 32 | 18 | 4.45 | 29 (74.36 %) |
| Ventral Tegmental Area (ext. MTT/pHyp) | 6 | -10 | -6 | 5.05 | 29 (74.36 %) |
| ventral part of the anterior Insula | 34 | 20 | -12 | 4.48 | 29 (74.36 %) |
| ventral part of the anterior Insula | -32 | 12 | -4 | 4.51* | 29 (74.36 %) |
| inferior part of the ventroanterior Thalamus | 4 | -10 | 4 | 4.88 | 28 (71.79 %) |
| inferior part of the ventroanterior Thalamus | -6 | -12 | 4 | 4.03* | 28 (71.79 %) |
| Ventral Tegmental Area (ext. MTT/pHyp) | -6 | -12 | -8 | 4.94 | 28 (71.79 %) |
| Caudal-Rostral Linear Raphe (ext. PTg) | -2 | -28 | -14 | 3.24 | 27 (69.23 %) |
| lateral Orbitofrontal Cortex (vlPFC) | 38 | 34 | -8 | 5.66* | 27 (69.23 %) |
| Lateral Geniculate Nucleus | -24 | -22 | -8 | 3.70 | 27 (69.23 %) |
| Lateral Geniculate Nucleus | 24 | -22 | -8 | 3.68 | 27 (69.23 %) |
| posterior Hippocampus | 24 | -38 | -2 | 5.63* | 27 (69.23 %) |
| lateral Amygdala (ext. aHIP) | -28 | -8 | -14 | 3.44 | 27 (69.23 %) |
| Caudal-Rostral Linear Raphe (ext. PTg) | 6 | -26 | -12 | 4.45 | 26 (66.67 %) |
| Subcoeruleus | 2 | -24 | -6 | 5.01 | 26 (66.67 %) |
| posterior Hippocampus | -26 | -40 | -2 | 4.35* | 26 (66.67 %) |
| Calcarine Sulcus | -8 | -62 | 18 | 3.18 | 26 (66.67 %) |
| Lateral Prefrontal Cortex | 26 | 48 | 4 | 3.73 | 25 (64.10 %) |
| Lateral Prefrontal Cortex | -24 | 54 | 10 | 3.8 | 25 (64.10 %) |
| lateral Orbitofrontal Cortex (vlPFC) | -36 | 32 | -10 | 3.48 | 25 (64.10 %) |
| lateral Amygdala (ext. aHIP) | 30 | -8 | -14 | 3.39 | 25 (64.10 %) |
| posterior MidCingulate Cortex | 2 | -14 | 40 | 2.93 | 25 (64.10 %) |
| dorsomedial Prefrontal Cortex | 4 | 38 | 38 | 2.65 | 25 (64.10 %) |
| Calcarine Sulcus | 12 | -54 | 16 | 3.05 | 25 (64.10 %) |
| *Note. Only peaks ≥ 60% are displayed. All peak were statistically significant in the One-Sample Permutation Test (TFCE FWE<0.05, 5,000 permutations). * top peak intensity was found in neighboring voxels.* MTT = Mammillothalamic Tract; pHyp = posterior Hypothalamus; PTg = Pedunculotegmental nucleus; vlPFC = ventrolateral Prefrontal Cortex; aHIP = anterior Hippocampus; | | | | | |

| **Supplementary Table 6.** Most replicable peaks of the Voxel-based Morphometry Network underpinning Trait Aggression | | | | | |
| --- | --- | --- | --- | --- | --- |
| Regions | MNI Coordinates | | | Peak Intensity (t-values) | Replicability (N, %) |
|  | x | y | z |  |  |
| dorsal part of the middle Insula | 34 | -2 | 10 | 7.73 | 23 (74.19 %) |
| posterior Insula | -42 | -12 | -4 | 9.23 | 22 (70.97 %) |
| dorsal part of the middle Insula | 34 | 34 | -8 | 4.4 | 21 (67.74 %) |
| posterior Insula | 36 | -12 | 8 | 8.73 | 21 (67.74 %) |
| medial Orbitofrontal Cortex | 2 | 40 | -12 | 6.63 | 21 (67.74 %) |
| Claustrum | -30 | 16 | 0 | 7.37 | 21 (67.74 %) |
| Superior Temporal Gyrus | 46 | -8 | -8 | 8.72 | 21 (67.74 %) |
| Claustrum | 30 | 16 | 0 | 7.07 | 20 (64.52 %) |
| Temporal Pole (ext. lAMY) | -30 | 4 | -24 | 7.27 | 20 (64.52 %) |
| Temporal Pole (ext. lAMY) | 30 | 8 | -26 | 6.75 | 20 (64.52 %) |
| posterior Midcingulate Cortex | 4 | -22 | 44 | 8.11 | 19 (61.29 %) |
| anterior Midcingulate Cortex | 2 | 24 | 36 | 7.13 | 19 (61.29 %) |
| *Note. Only peaks ≥ 60% are displayed. All peak were statistically significant in the One-Sample Permutation Test (TFCE FWE<0.05, 5,000 permutations). * top peak intensity was found in neighboring voxels.* lAMY = lateral Amygdala | | | | | |

| **Supplementary Table 7.** Gene-Set Enrichment Analysis of GO Biological Processes (Enrichr) | | | | | |
| --- | --- | --- | --- | --- | --- |
| Term | Gene overlap | P-value | Adjusted P-value | Odds Ratio | Combined Score |
| Cell-Cell Adhesion via Plasma-Membrane Adhesion Molecules (GO:0098742) | 9 out of 171 | 1.38E-06 | 0.00126541 | 9.36 | 126.28 |
| Synaptic Membrane Adhesion (GO:0099560) | 4 out of 17 | 3.35E-06 | 0.00153446 | 50.09 | 631.41 |
| Regulation of Neuron Projection Development (GO:0010975) | 8 out of 176 | 1.59E-05 | 0.00485962 | 7.95 | 87.85 |
| Nervous System Development (GO:0007399) | 11 out of 480 | 2.31E-04 | 0.04640227 | 3.96 | 33.14 |
| Negative Regulation of Cell Projection Organization (GO:0031345) | 4 out of 51 | 2.98E-04 | 0.04640227 | 13.83 | 112.27 |
| Synapse Organization (GO:0050808) | 6 out of 144 | 3.04E-04 | 0.04640227 | 7.15 | 57.90 |
| *Note.* Only biological processes with statistical significance of p<0.05 after adjustments are displayed. | | | | | |

| **Supplementary Table 8.** Functions of the 5 genes most strongly expressed in aggression-related brain networks | | |
| --- | --- | --- |
| **Gene** | **Full Name** | **Primary Function** |
| **SYT10** | Synaptotagmin 10 | Mediates calcium-dependent exocytosis; involved in neuroprotection and vesicle release. |
| **TENM3** | Teneurin Transmembrane Protein 3 | Guides axon development and synaptic organization; associated with eye development abnormalities. |
| **NAV3** | Neuron Navigator 3 | Regulates axonal growth and cytoskeletal dynamics during neuronal development. |
| **SCN9A** | Sodium Voltage-Gated Channel Alpha Subunit 9 | Encodes Nav1.7 sodium channel; crucial for pain signaling and sensory neuron excitability. |
| **ANKRD50** | Ankyrin Repeat Domain 50 | Participates in endosomal protein sorting; interacts with retromer complex for membrane recycling. |

### **Supplementary Figures**

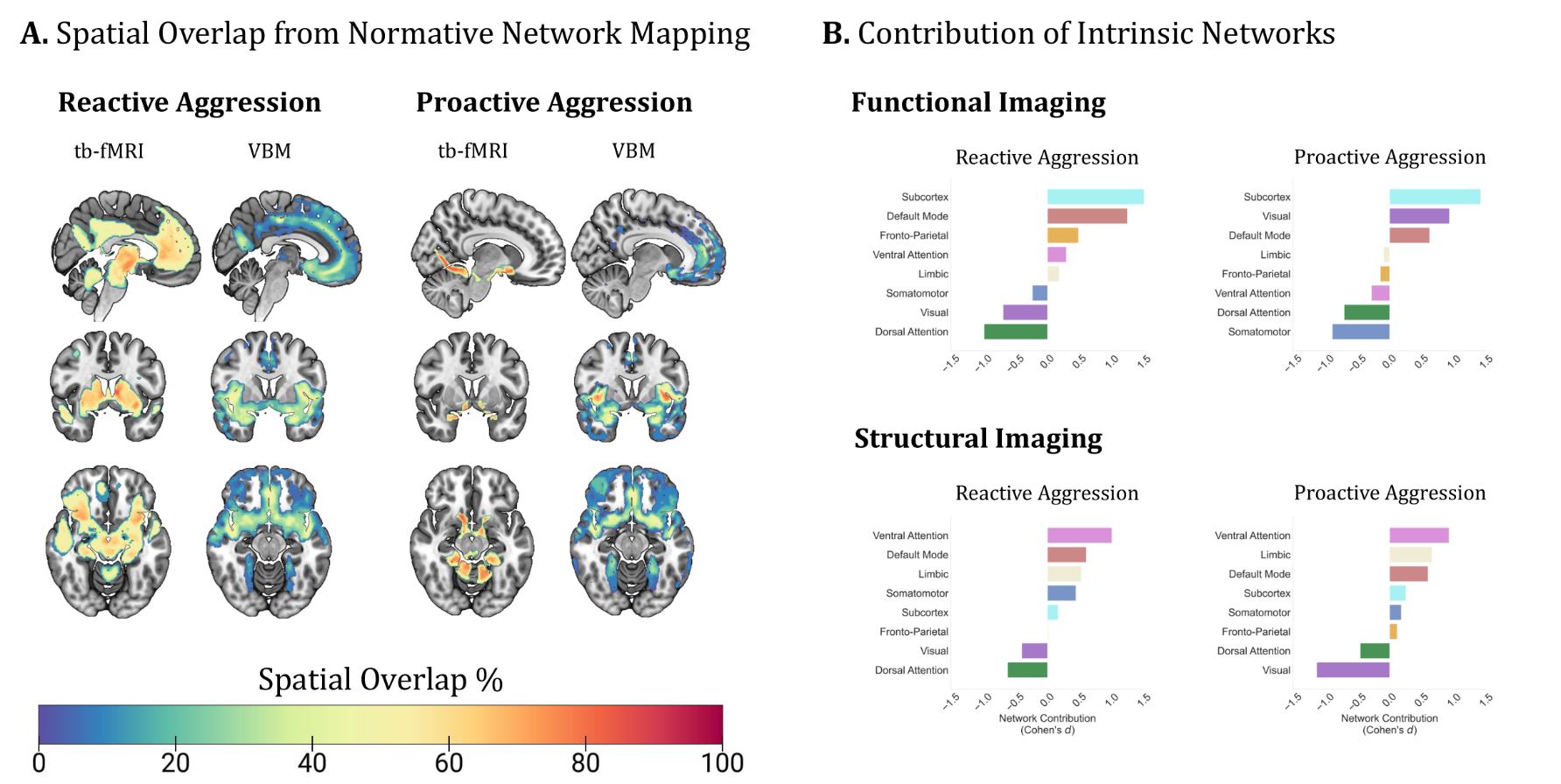

**Supplementary Fig. 1.** Functional and Structural Brain Networks Underpinning Reactive and Proactive Aggression. Poor replicability in traditional coordinate-based meta-analysis (ALE algorithm, Dugré Dugré) was detected across both reactive (*Functional* < 10.81% overlap; *Structural* < 13.33% overlap) and proactive aggression (*Functional* < 15.79% overlap, *Structural* < 18.18 % overlap). **Panel A.** Strong network replicability was also found across both reactive (*Functional* < 86.49% overlap; *Structural* < 60 % overlap) and proactive aggression (*Functional* < 89.47% overlap; *Structural* < 90.91% overlap). **Panel B.** Functional Brain Networks underpinning Reactive Aggression was mainly characterized by connectivity of the Subcortex (*d* = 1.59), default mode (*d* = 1.24), and frontoparietal networks (*d* = 0.48) to a lesser extent, while Proactive aggression was mainly characterized by connectivity of the Subcortex (*d* = 1.59), visual (*d* = 0.93), default mode (*d* = 0.62) networks. Structural Brain Networks underpinning Reactive Aggression was mainly characterized by connectivity of the Ventral attention (*d* = 0.99), default mode (*d* = 0.60), limbic (*d* = 0.53), Somatomotor (*d* = 0.44), while Proactive aggression was mainly characterized by connectivity of the Ventral attention (*d* = 0.92), limbic (*d* = 0.66), default mode (*d* = 0.60) networks. Tb-fMRI = Task-based functional magnetic resonance imaging; VBM = Voxel-based Morphometry Studies.

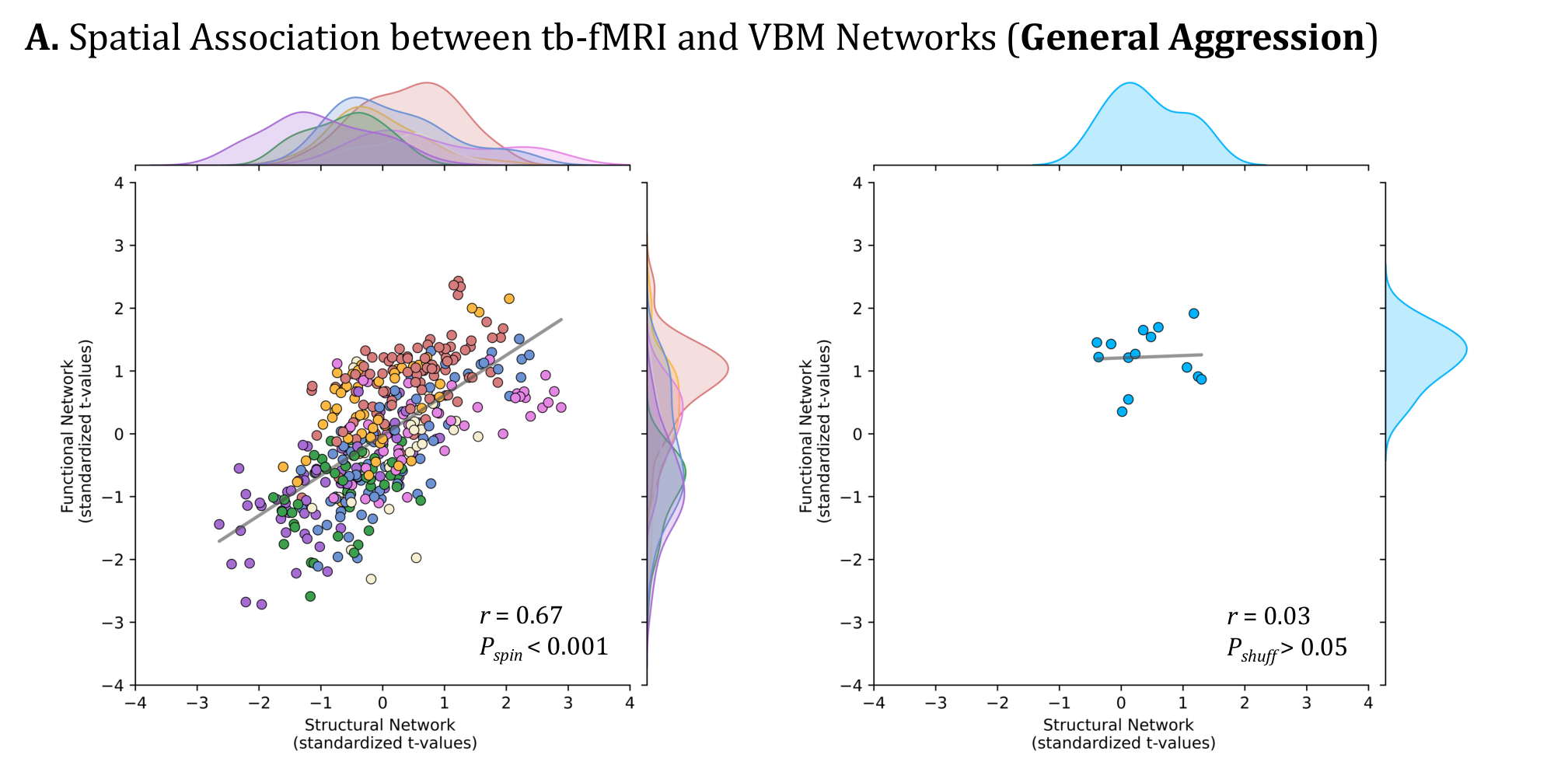

**Supplementary Fig. 2.** Spatial correlation between Functional and Structural Brain Networks underpinning **General Aggression**. Left Panel represents correlation between t-values of the 400 cortical regions from the functional brain network and those of from the structural brain networks. Statistical significance was determined via spin test with 5,000 permutations. Right Panel represent the correlation between t-values of the 14 subcortical regions from the functional brain network and those from the structural brain networks. Statistical significance was determined via shuffle test with 5,000 permutations.

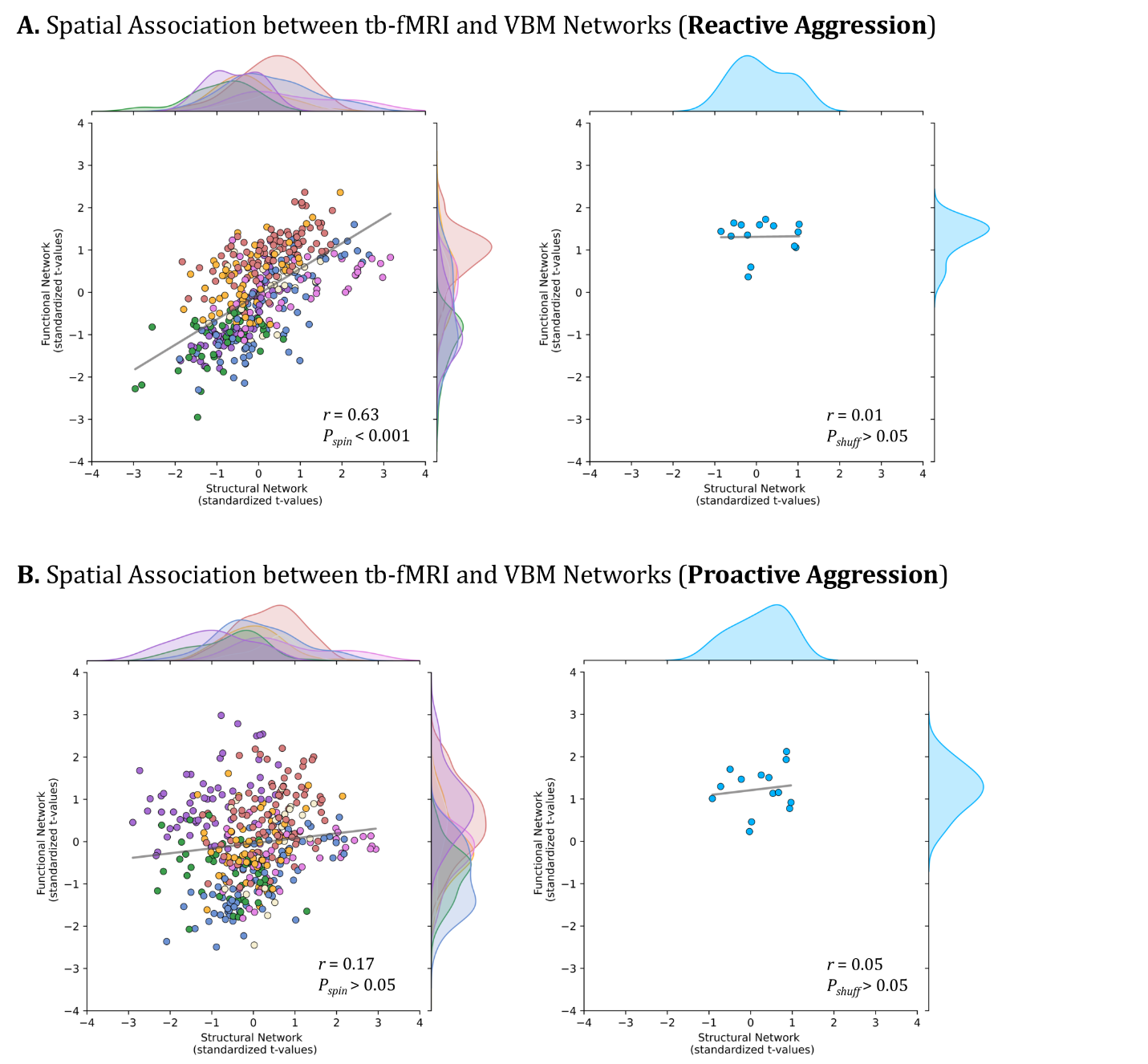

**Supplementary Fig. 3.** Spatial correlation between Functional and Structural Brain Networks underpinning subtypes of Aggression. **A.** Reactive Aggression. **B.** Proactive Aggression. Left Panels represents correlation between t-values of the 400 cortical regions from the functional brain network and those of from the structural brain networks. Statistical significance was determined via spin test with 5,000 permutations. Right Panels represent the correlation between t-values of the 14 subcortical regions from the functional brain network and those from the structural brain networks. Statistical significance was determined via shuffle test with 5,000 permutations.

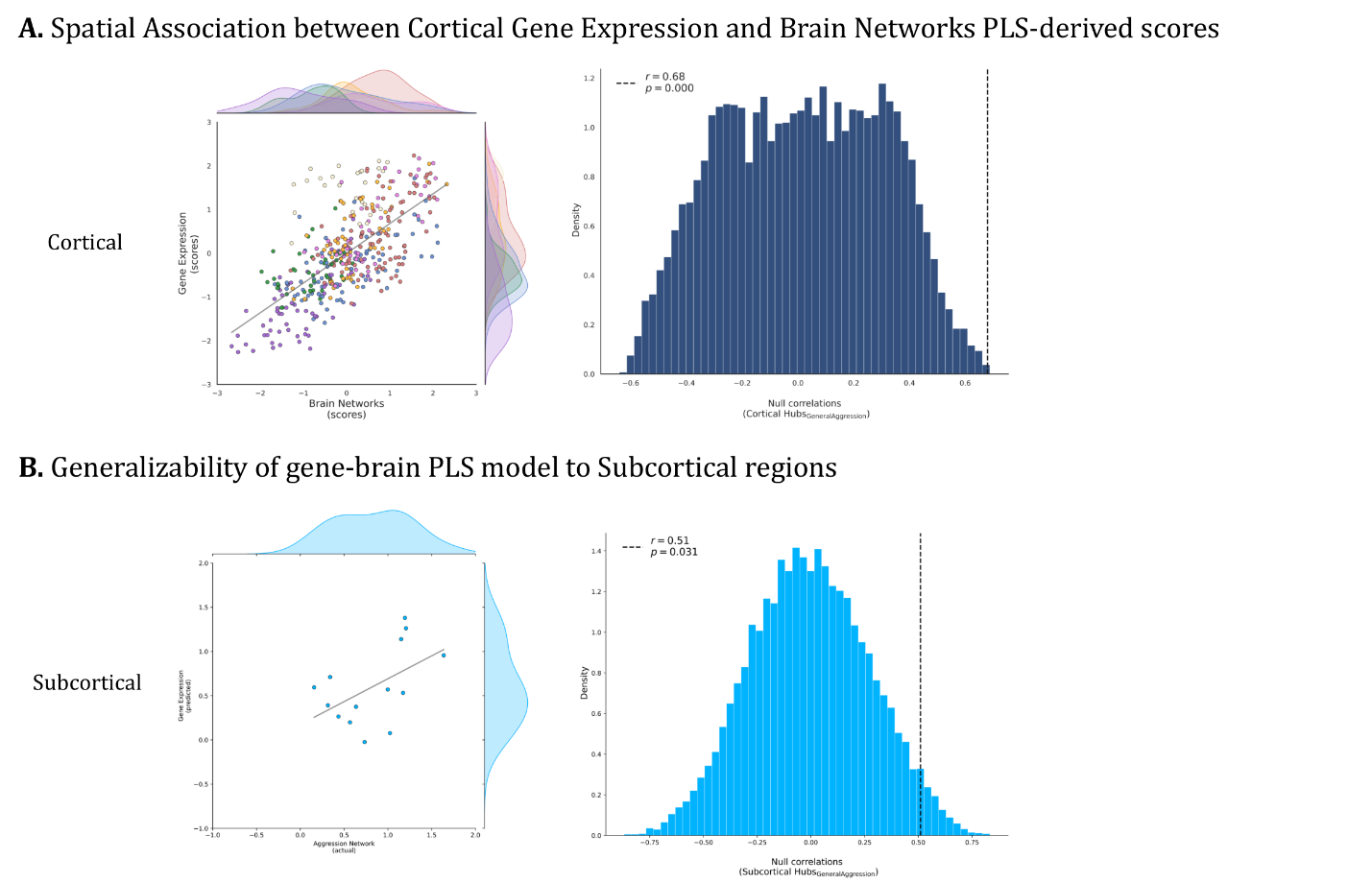

**Supplementary Fig. 4.** Partial Least Square Analysis Findings. **A.** Spatial correlation between projected cortical gene expression and aggression brain networks scores derived from the partial least square analysis. Left Panel depicts the correlation between gene expression and imaging composite scores of the PLS-derived latent variables across 400 cortical regions. Right Panel reflect the statistical significance of the gene-brain association, determined via spin test with 5,000 permutations. **B.** Generalizability of the model across subcortical regions was investigated through shuffle permutation test. Left Panel depicts the correlation between gene expression and imaging composite scores of the PLS-derived latent variables across 14 subcortical regions. Right Panel reflect the statistical significance of the gene-brain association, determined via shuffle test with 5,000 permutations.

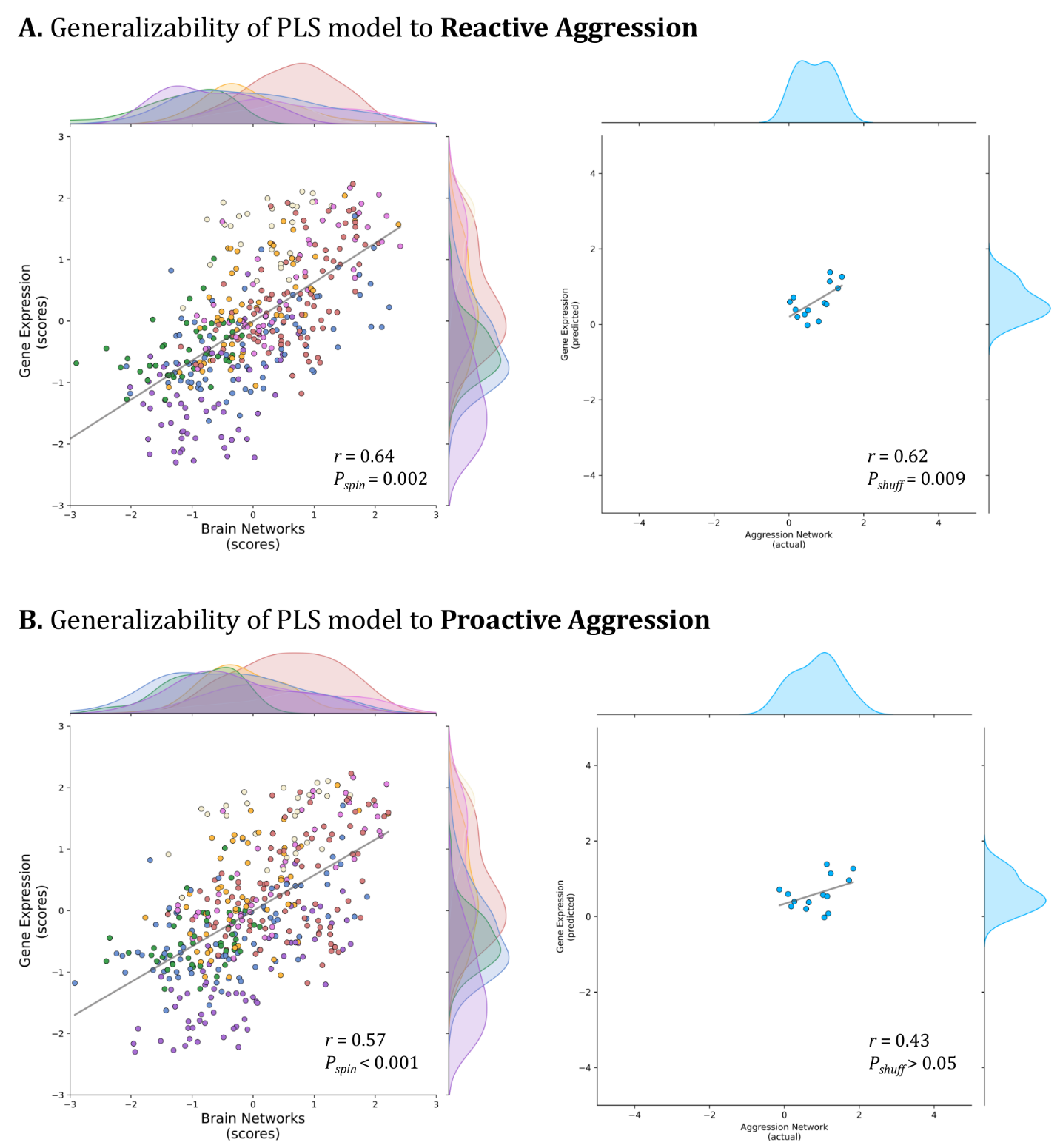

**Supplementary Fig. 5.** Generalizability of the PLS model to Subtypes of Aggression. **A.** Reactive Aggression Brain Networks. **B.** Generalizability of the PLS model to **Proactive Aggression** Brain Networks. Left Panels depict the correlation between gene expression and imaging composite scores of the PLS-derived latent variables across 400 cortical regions. Statistical significance was determined via spin test with 5,000 permutations. Right Panels depict the correlation between gene expression and imaging composite scores of the PLS-derived latent variables across 14 subcortical regions. Statistical significance was determined via shuffle test with 5,000 permutations.
